# The basis of antigenic operon fragmentation in *Bacteroidota* and commensalism

**DOI:** 10.1101/2023.06.02.543472

**Authors:** Nicholas C. Bank, Vaidhvi Singh, Brandon Grubb, Blake McCourt, Aaron Burberry, Kyle D. Roberts, Alex Rodriguez-Palacios

## Abstract

The causes for variability of pro-inflammatory surface antigens that affect gut commensal/opportunistic dualism within the phylum *Bacteroidota* remain unclear (1, 2). Using the classical lipopolysaccharide/O-antigen ‘*rfb* operon’ in *Enterobacteriaceae* as a surface antigen model (5-gene-cluster *rfbABCDX*), and a recent *rfbA-*typing strategy for strain classification (3), we characterized the architecture/conservancy of the entire *rfb* operon in *Bacteroidota*. Analyzing complete genomes, we discovered that most *Bacteroidota* have the *rfb* operon fragmented into non-random gene-singlets and/or doublets/triplets, termed ‘minioperons’. To reflect global operon integrity, duplication, and fragmentation principles, we propose a five-category (infra/supernumerary) cataloguing system and a Global Operon Profiling System for bacteria. Mechanistically, genomic sequence analyses revealed that operon fragmentation is driven by intra-operon insertions of predominantly *Bacteroides*-DNA (*thetaiotaomicron/fragilis*) and likely natural selection in specific micro-niches. *Bacteroides*-insertions, also detected in other antigenic operons (fimbriae), but not in operons deemed essential (ribosomal), could explain why *Bacteroidota* have fewer KEGG-pathways despite large genomes (4). DNA insertions overrepresenting DNA-exchange-avid species, impact functional metagenomics by inflating gene-based pathway inference and overestimating ‘extra-species’ abundance. Using bacteria from inflammatory gut-wall cavernous micro-tracts (CavFT) in Crohn’s Disease (5), we illustrate that bacteria with supernumerary-fragmented operons cannot produce O-antigen, and that commensal/CavFT *Bacteroidota* stimulate macrophages with lower potency than *Enterobacteriaceae*, and do not induce peritonitis in mice. The impact of ‘foreign-DNA’ insertions on pro-inflammatory operons, metagenomics, and commensalism offers potential for novel diagnostics and therapeutics.

## Introduction

An operon is a functional unit of DNA that consists of a cluster of contiguous genes, transcribed together to control cell functions, including the production of the O-antigen component of lipopolysaccharides (LPS), which is widely present and variable in gram-negative bacteria. The phylum *Bacteroidota* (*Bacteroidetes*), composed primarily of gram-negative gut commensals (6-10), is also known to have several opportunistic pathogenic species (‘pathobionts’) (11-19), which for unclear reasons have commensal/pathogenic dualism. Concerningly, species from the phylum have been proposed as future probiotics because some strains modulate gut immunity locally (6, 20, 21) or influence the susceptibility to chronic extra-intestinal diseases (*e*.*g*., *Parabacteroides goldsteinii* attenuates obstructive pulmonary disease (15).

The precise role of *Bacteroidota* in chronic inflammatory bowel diseases (IBD), namely Crohn’s disease (CD), remains undefined (22). Supporting a pathogenic role in IBD complications, we recently discovered that the inflamed bowel of patients with surgical/severe CD have cavitating ‘cavernous fistulous tract’ micropathologies (CavFT) resembling cavern formations, harboring cultivable bacteria, including *Escherichia coli* and *Bacteroidota* (5, 23). Focused on a few CavFT species, genomic analyses of consecutive *Bacteroidota* isolates (*Parabacteroides distasonis*) from unrelated patients that underwent surgery for CD suggested, for the first time, that certain bacteria (from a novel lineage in NCBI databases) are adapting to CavFTs, swapping large fragments of DNA with *Bacteroides*, and are likely transmissible in the community (23). To classify *P. distasonis* and other cultivable *Bacteroidota*, and to facilitate the orderly study of such commensal/pathogenic dualism in the phylum, we recently proposed the use of the *rfbA* gene for genotyping *Bacteroidota*. Of interest, *rfbA*-typing suggested that historical *P. distasonis* strains isolated from pathological sources belonged to one of four *rfbA* types (3).

Compared to the lipid A gene *lpxK* (which was highly conserved), *rfbA*-typing studies demonstrated that the O-antigen (*rfb*) genes are sufficiently variable to be better associated with the variable pathogenic potential of *Bacteroidota* for bacterial genotypic classification. Since lipid A in *Bacteroidota* induces lower TLR4 inflammatory activation (6, 15, 21, 24) compared to *E. coli*, and since the role of O-antigen/LPS in *Bacteroidota* is poorly understood (20), herein, we conducted an expanded typing analysis across all genes of the *rfb* operon in *Bacteroidota* using *i)* existing complete genomes and NCBI genome databases, *ii)* new genomes sequenced from CavFT, *iii)* the classical *Enterobacteriaceae rfb* operon contiguity as referent, and *vi)* using *Parabacteroides* and *Alistipes* as emerging pathogenic/probiotic models (1, 2) to identify potential genomic features impacting surface antigens favoring *Bacteroidota* commensal/pathobiont dualism.

## Results

### The classic *rfb* operon is contiguous in *Enterobacteriaceae*

As a referent to illustrate the arrangement of *rfb* genes involved in the O-antigen synthesis operon in *Enterobacteriaceae*, we analyzed reference genomes from *E. coli, Klebsiella variicola*, and *Salmonella enterica*. A schematic of the LPS/O-antigen molecule is shown in **Figure 1A**. In *E. coli* K12, **Figure 1B** illustrates the contiguous arrangement of five genes which functionally enable the *en bloc* transcription of the *rfb* operon (*rfbABCDX*). Regardless of transcriptional orientation (positive/negative sense), other *Enterobacteriaceae* have the same contiguous arrangement of the *rfb* genes. Notably, while the length of *rfb* operons vary across this family, bacteria predominantly have at least four genes (*rfbABCD)*, organized contiguously, with sporadic singly duplicated genes. Contiguity is so conserved within the family that *Salmonella* spp. have operons with up to 15 consecutive *rfb* genes (**Supplementary Figure 1A-B**).

**Figure 1.**
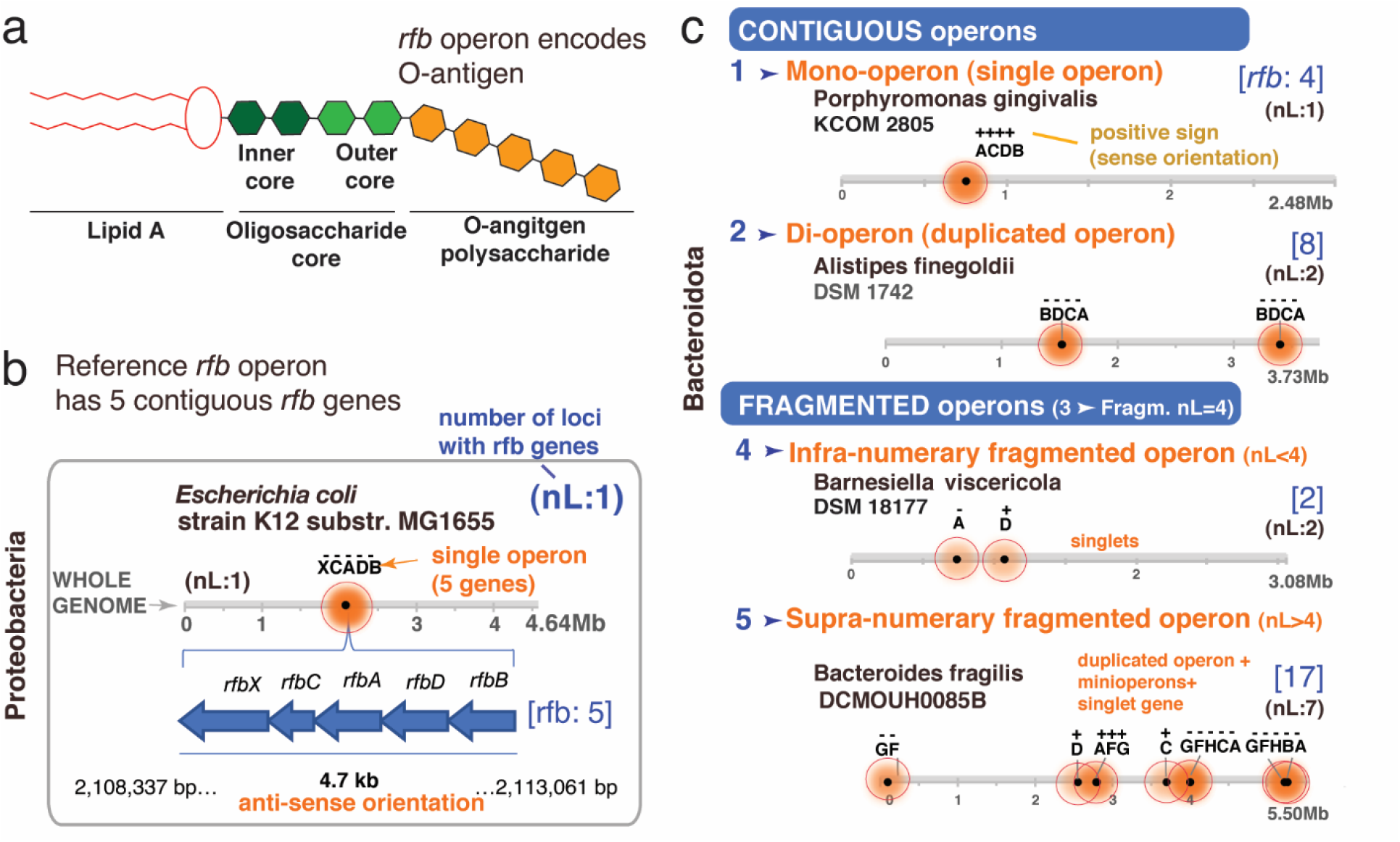
The *rfb* operon in *Enterobacteriaceae* is contiguous, but in *Bacteroidota* it is often fragmented into ‘minioperon’. **A)** Lipopolysaccharide (LPS) and O-antigen schematics. **B)** Classical *rfb* operon with 5 contiguous genes in *E. coli* K12 (*rfbXABCD;* XCADB; [*rfb*:5]). Horizontal lines represent the bacterial genome length, distribution of *rfb* genes/operons (shaded circles; darker = more genes) and *rfb* gene orientation (+, sense; -, antisense). nL: n of *rfb* clusters or gene singlet loci. Note orientations/duplications. **C)** Five categories of operon arrangement & ‘minioperon’ fragmentation in *Bacteroidota*. **Supplementary Figure 1C** illustrates in context *rfb* operons/fragmentation for *Alistipes, Bacteroides, Parabacteroides, Prevotella, Paraprevotella, Barnesiella, Tannerella, Odoribacter* and *Porphyromonas*.

### The *rfb* operon in *Bacteroidota* is often fragmented into ‘minioperons’

To assess the spatial integrity of the *rfb* operon in *Bacteroidota*, we next examined the *rfb* gene profile (presence/absence) in available complete genomes. Analysis revealed that the *rfb* genes in *Parabacteroides, Bacteroides*, and *Prevotella* were not contiguous but fragmented and dispersed throughout the genome. In contrast, *Alistipes, Porphyromonas* and *Odoribacter* have *rfb* operons composed of the same primordial genes as *Enterobacteriaceae (rfbACDB)*, being intact and/or duplicated, supporting their genomic potential to be functionally capable of LPS-proinflammatory induction. Within the phylum, operon fragmentation products, herein referred to as *rfb* “minioperons”, result in various combinations of *rfb* singlet genes and doublets/triplets with variable orientations (sense: +; antisense: -, **Supplementary Figure 1C**). With *E. coli* as referent, using four *rfb* genes (n=4) as the median maximum number of contiguous genes observed among *Bacteroidota* in this study, herein we propose that *rfb* operons in *Bacteroidota* can be classified into at least five categories: *i)* Mono-operon (single contiguous operon, regardless of numbers of genes in cluster/locus), *ii)* Di-operon (duplicated operon), *iii)* fragmented normo-numerary operon (operon fragmented with 4 *rfb* clusters/loci, nL=4), *iv)* fragmented infra-numerary operon (nL<4 clusters/loci), and *v)* fragmented supernumerary operon (nL>4 clusters/loci; **Figure 1C**). Used alongside a baseline referent, this system of categorization and cataloguing may be applied to any operon system.

Envisioning the potential combinatorial variability for the O-antigen via the theoretical pairwise combination of *rfb* genes in *Bacteroidota*, we next determined that mathematical permutations of minioperons could yield thousands of possible combinations unlikely to yield functional O-antigen polysaccharides (**Supplemental Figure 2A**), which proves technically challenging to verify experimentally for all bacteria by requiring innumerable grow conditions.

### Minioperon patterns in *Parabacteroides* suggests *rfb* gene distancing mechanism

Since the study of potential operon fragmentation mechanisms could be better achieved using strains with fully sequenced genomes, we next conducted an arrangement analysis to determine if *rfb* operon fragmentation was common within any given species and if they followed statistically significant patterns among unrelated strains of the same species. Using *P. distasonis* as a model *Bacteroidota* species and strain ATCC8503 (peritonitis, USA/1935) as the referent species for the genus *Parabacteroides*, analyses revealed that *P. distasonis* have their *rfb* operons invariably fragmented (12/12, Fisher’s exact P<0.00001), following unique patterns of gene combinations, duplications, or orientations that are significantly more likely to be supernumerary than infra-numerary (Fisher’s exact P=0.0001, **Figures 2A** and **1C**). These findings suggest that some species are more likely to gain *rfb* loci (*P. distasonis*), compared to others which could be more likely to lose *rfb* loci (*Barnesiella viscericola*).

**Figure 2.**
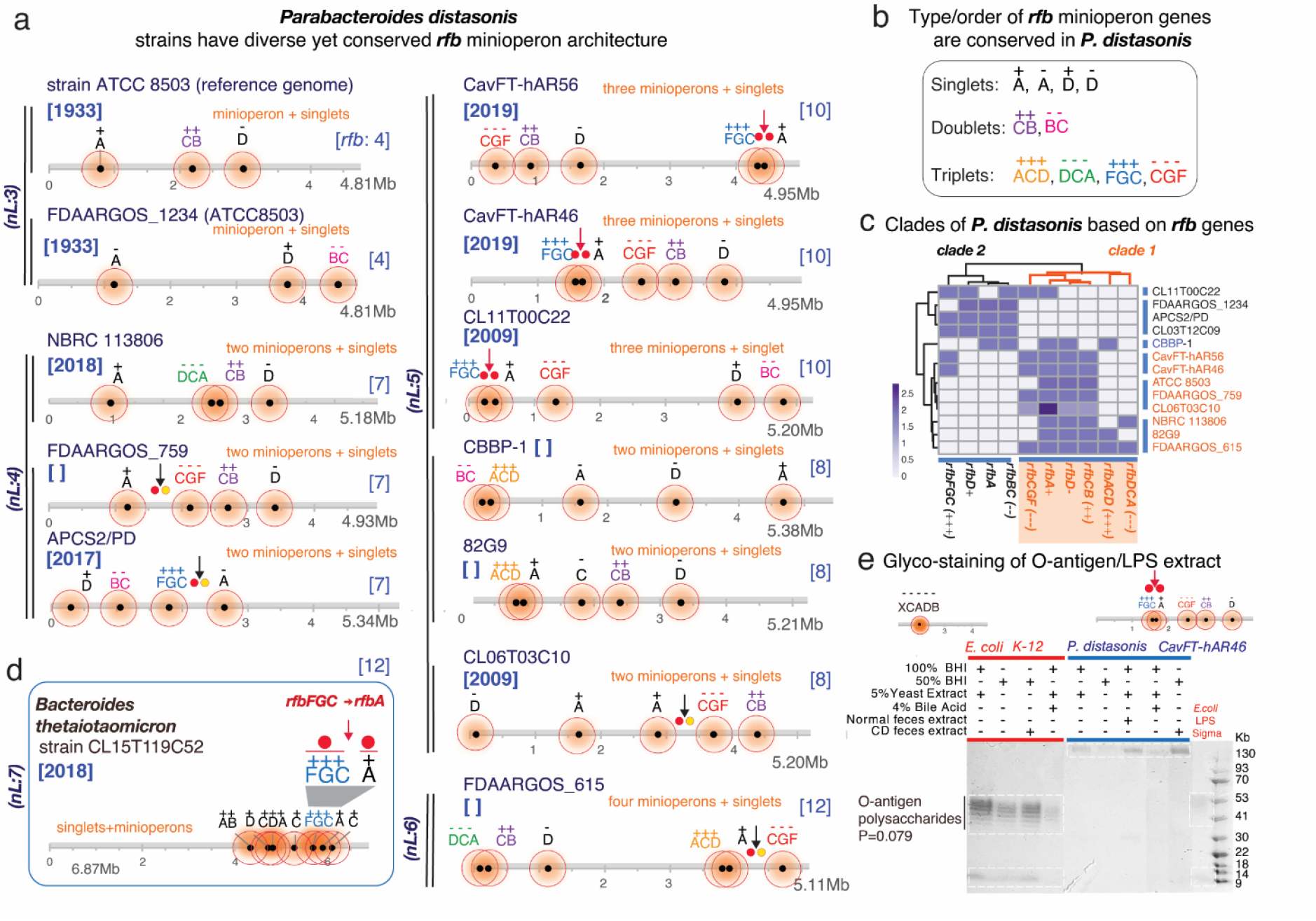
The occurrence, patterns and inversions of *rfb* minioperons in *P. distasonis* and *B. thetaiotaomicron* suggests mechanism of operon fragmentation in *Bacteroidota*. **A)** Fragmentation of *rfb* operon *in P. distasonis* is in modern times supernumerary compared to ATCC8503 strain from USA/1933. **B)** Fragmentation has resulted in conserved singlet, doublet/triplet patterns in *P. distasonis*. **C)** Heatmap clustering of *P. distasonis* strains based on *rfb* minioperon shows distinct clades. Additional information is available in **Supplementary Figure 2B. D**) Unique *rfbFGC->rfbA* minioperon distancing pattern (downward red/black arrows and solid circles) in *B. thetaiotaomicron* is also present in novel CavFT strains of *P. distasonis* from gut wall lesions in Crohn’s disease. Notice the orientation sense and patterns. **E**) SDS-PAGE of LPS extract analysis of *E. coli* and *P. distasonis* CavFT-46 shows that *P. distasonis* is unable to produce O-antigen polysaccharides in diverse growth conditions (Fisher’s exact P=0.079).

Fragmentation patterns were also highly conserved among *P. distasonis*. Specifically, *rfbA* and *rfbD* genes were present in minioperons or as singlets, whereas the genes *rfbB, rfbC, rfbG* and *rfbF* were exclusively present only as doublet- or triplet-minioperon arrangements (**Figure 2B**). Based on such conservancy, *P. distasonis* strains belong to at least two distinctive phylogenetic clades (**Figure 2C, Supplementary Figure 2B**), suggesting that not all *P. distasonis* would be the same, and emphasizing the need to better classify *Bacteroidota* isolates for functional studies.

Mechanistically insightful, outside of this species we found that a unique combination and distancing of two *rfb* minioperons (*rfbFGC->rfbA*; abbreviation for *rfbFGC*(+++) positive-sense triplet followed by a *rfbA*(+) positive-sense singlet), overrepresented in some *P. distasonis* strains (n=6/12, Fisher’s exact P<0.001, vs. numerous other possibilities), was also present in a *B. thetaiotaomicron* strain CL15T119C52 (**Figure 2D**), a phenomenon not previously reported in the literature.

Considering that *B. thetaiotaomicron* lacks LPS-polysaccharide formation (25) and has anti-inflammatory and immunomodulatory properties (18, 19), the presence of such *rfbFGC->rfbA* fragmentation pattern in both *P. distasonis/B thetaiotaomicron* suggests a mechanism for operon fragmentation that could explain beneficial effects for both species/strains (2). Remarkably, an inter-genus *rfbFGC->rfbA* pattern, exclusively present in *P. distasonis* strains of CavFT origin and CL11T00C22 (5), have conserved minioperon sequences but variable inter-minioperon *rfbFGC->rfbA* distances, suggesting that DNA insertion within the flanking *rfb* genes could be the reason for gene-gene separation in an ongoing permissible process of gene exchange between *P. distasonis* and *B thetaiotaomicron*.

Of novelty, variable inter-minioperon orientations for the same *rfbFGC->rfbA* pattern in other *P. distasonis* genomes (APCS2/PD, FDAARGOS_615, and CL06T03C10) suggest the existence of conserved *rfbFGC->rfbA* inversions with a pivot point in the inter-minioperon *rfbFGC->rfbA* region. This observation indicates that such a potential gene-gene separation mechanism is highly conserved, yet variable. Furthermore, it offers an evolutionary explanation to the presence of unique arrangements across certain strains, lineages, or possible niches that could reflect favored co-habitation of both genera, genetic exchange, selection, and niche adaptation (notice the red vs. black arrows for *rfbFGC->rfbA* in **Figures 2A & 2D**).

To elucidate the effect of minioperons on LPS/O-antigen production and structure, we opted to verify the presence/absence of the O-antigen polysaccharide in *P. distasonis* CavfT-hAR46, under five unique growth conditions, as a cultivable *Bacteroidota* model with 5 *rfb* minioperon fragments (vs. *E. coli* with one *rfb* operon). Of functional importance, the classical *rfb* operon in *E. coli* effectively produced a typical O-antigen/polysaccharide as expected in all conditions, with repeated bands between 41 and 53 kb in SDS-PAGE (**Figure 2E**) (26, 27). However, as expected for a *P. distasonis* strain with supernumerary *rfb* fragmentation, *P. distasonis* did not yield O-antigen polysaccharides in any of the conditions, suggesting that supernumerary *rfb* fragmentations are likely to be non-functional (5/5 vs. 0/5, Fisher’s exact P=0.0079). As a result, lipooligosaccharide products (LOS) were produced instead of LPS, similar to what has recently been described for *B. thetaiotaomicron* (25).

### The *rfb* operon in *Alistipes* is intact or duplicated suggesting benefit for survival

Contrasting *Parabacteroides*, the *Alistipes* genus primarily exhibits no operon fragmentation. Unique conserved organization and orientation was observed in all examined *Alistipes* species which have 4-gene contiguous *rfb* operons (*BDCA* or *CADB;* **Figure 3A**). When fragmentation was present (*A. shahii, A. dispar*), *Alistipes* have minioperon doublets (*rfbCA, DB, AC, GF*) not seen in *Parabacteroides*, furthers supporting that genus-specific mechanisms of operon fragmentation or conservation vary across *Bacteroidota*.

**Figure 3.**
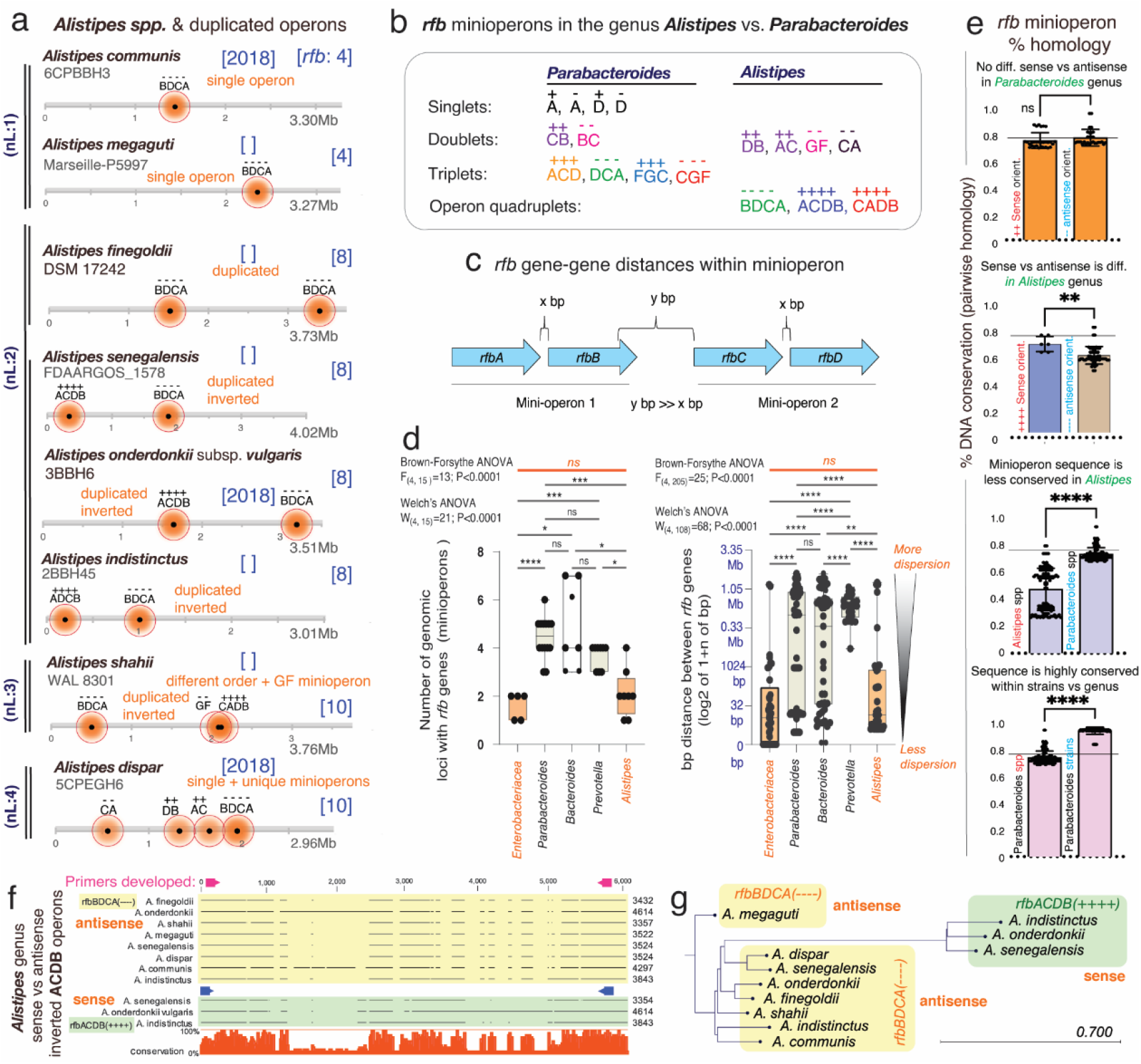
The *rfb* operon in *Alistipes* is contiguous and duplicated suggesting evolutionary benefit. **A)** *Alistipes* has contiguous *rfb* operons with frequent duplication and less common incorporation of *rfb* minioperon duplets. **B)** Patterns of conserved minioperons in *Alistipes* differ from *Parabacteroides*. **C)** Schematics of gene-gene *rfb* distances measured within and between minioperons. **D)** *Parabacteroides* and *Bacteroides* demonstrate the greatest variance in number of *rfb* operon fragments and gene-gene distances. *Alistipes* and *Enterobacteriaceae* are similarly contiguous. Intergene distances for *Parabacteroides, Bacteroides* and *Prevotella* were greater than *Enterobacteriaceae* (0.54±0.84Mb, 0.49±0.86Mb, 0.34±0.32Mb, respectively, vs. 0.13±0.46Mb, P<0.001). *Alistipes* gene distribution is similar to *Enterobacteriaceae* (0.17±0.49Mb, P=0.79). **E**) Minioperon sequence homologies for *Alistipes* vs. *Parabacteroides* based on minioperon orientation between and within genera/species. Two tailed-T tests P<0.01 **, P<0.001 ***. **F)** Alignment and **G)** phylogeny based on *rfb* operon sequences. Note that *Alistipes* clusters are driven by the sense/antisense orientation of the operons.

Intriguingly, as a major difference within other *Bacteroidota*, most *Alistipes* species have duplicated operons (5/8 vs. 1/22, *B. fragilis*, Fisher’s exact P=0.0018), indicating that the genus has genomic potential for enhanced O-antigen production and possibly pro-inflammatory LPS effects that could be necessary or beneficial for *Alistipes* adaptation/survival. Alternatively, operon duplication with minimal fragmentation suggests that *i)* fragmentation is non-sustainable or rather deleterious for *Alistipes*, and/or *ii)* its genetic mechanisms of operon maintenance allow for operon variability (duplication, inversions, sequence), but not for gene-operon separation, unlike *Parabacteroides* in which fragmentation predominates (12/12 vs. 2/8, *A. shahii & A. dispar*, Fisher’s exact P=0.0049). No *rfb* singlets were observed in *Alistipes*.

The *rfb* minioperon types in *Alistipes* and *Parabacteroides* indicate that gene patterns and orientations are unique and vary, being potentially useful as signatures for lineage identification and classification (**Figure 3B**). When we quantified the magnitude of fragmentation (gene-gene distances in number of nucleotides) across genera, we found that *Parabacteroides, Bacteroides* and *Prevotella* had similarly more fragmentation on average than *Enterobacteriaceae* (P <0.001, P=0.0151, P=0.010, respectively), but not *Alistipes* compared to *Enterobacteriaceae* (2.1 vs. 1.6, P=0.89, **Figure 3C-D**).

### Minioperon conservation in *P. distasonis* encompasses 85 years of history

Using *Parabacteroides* and *Alistipes* as genus models for *Bacteroidota* (1, 2), we then quantified the *rfb* sequence conservation (% similarity among strains)(3), since conservation indicates functional/evolutionary advantages (28). Analysis showed that, irrespective of orientation, the doublet *rfbCB* sequences, present in 80% of *P. distasonis*, and the *rfbACD* triplets have homologies ranging from 72 to 99.8% (77.5±5.15%), contrasting the much lower operon similarities across *Alistipes* (range 27.5 to 83.7%; 50±16.4%, P<0.001; **Supplementary Figure 3**).

Considering that the *P. distasonis* strains examined in this study span 85 years and various geographic locations, from ATCC8503 (USA, c.1933) to 82G9 (South Korea, c.2018), the high minioperon sequence similarity (P<0.001 **Figure 3E**) indicates well-conserved genomic features across time and space. However, this conservation cannot explain the observed *rfbFGC->rfbA* variability. Instead, it implies that an independent genomic mechanism might be responsible for generating modern *rfbFGC->rfbA* variants over time, which were not present in the founding strain ATCC8503, 85 years ago.

Although the operons in *Alistipes* are more similarly organized (*BDCA*) than *Parabacteroides*, the sequences are more variable, depending on operon orientation (**Figure 3F-G**). Of interest, duplicated minioperons in *Parabacteroides* are virtually identical, which contrasts the sequence dissimilarity seen among duplicated *rfb* operons in *Alistipes* (T-Test P=0.039, **Figure 3E**). The variable homologies across *Alistipes* indicate that they may *i)* produce a more variable array of LPS/O-antigens, with at least two different types of O-antigens if operons are duplicated, and/or *ii)* have higher virulence *in vivo*. Given the emergence of *Alistipes* as an emerging genus in human diseases (1), we developed PCR primers for *rfbBDCA/ACDB* operons to facilitate their future study in *Alistipes* (**Supplementary Table 1**).

### Genome *‘rfb* operon profiling’ shows minioperon occurrence is not random

To further characterize the *rfb* genes and the genome-wide operon patterns in *Bacteroidota*, we propose a global ‘*rfb* operon profiling’ system (GOPS). Using our *rfbA*-typing methodology (3) and *P. distasonis* strains as a model, we showed that it is possible to discern the strains examined based on distinct genotypes for each *rfb* gene (*rfbB*-types n=2, *rfbC*-types n=6, *rfbD*-types n=4, *rfbF*-types n=2, *rfbG*-types n=2, **Figure 4A & Supplementary Figure 4**). By applying the previous scheme to generate a combined alphabetic-numeric profile of the entire *rfb* operon, accounting for both copy number and *rfb* genotype, we generated summary profiles for testing **(Figure 4B)**. Of note, this system may be broadly applied to type other operon genes. Remarkably, the GOPS profiles identified were determined to be reproducible, favoring profiles that are statistically different from random profiles, supporting the assumption that minioperon arrangements have been selected over time (**Figure 4C**).

**Figure 4.**
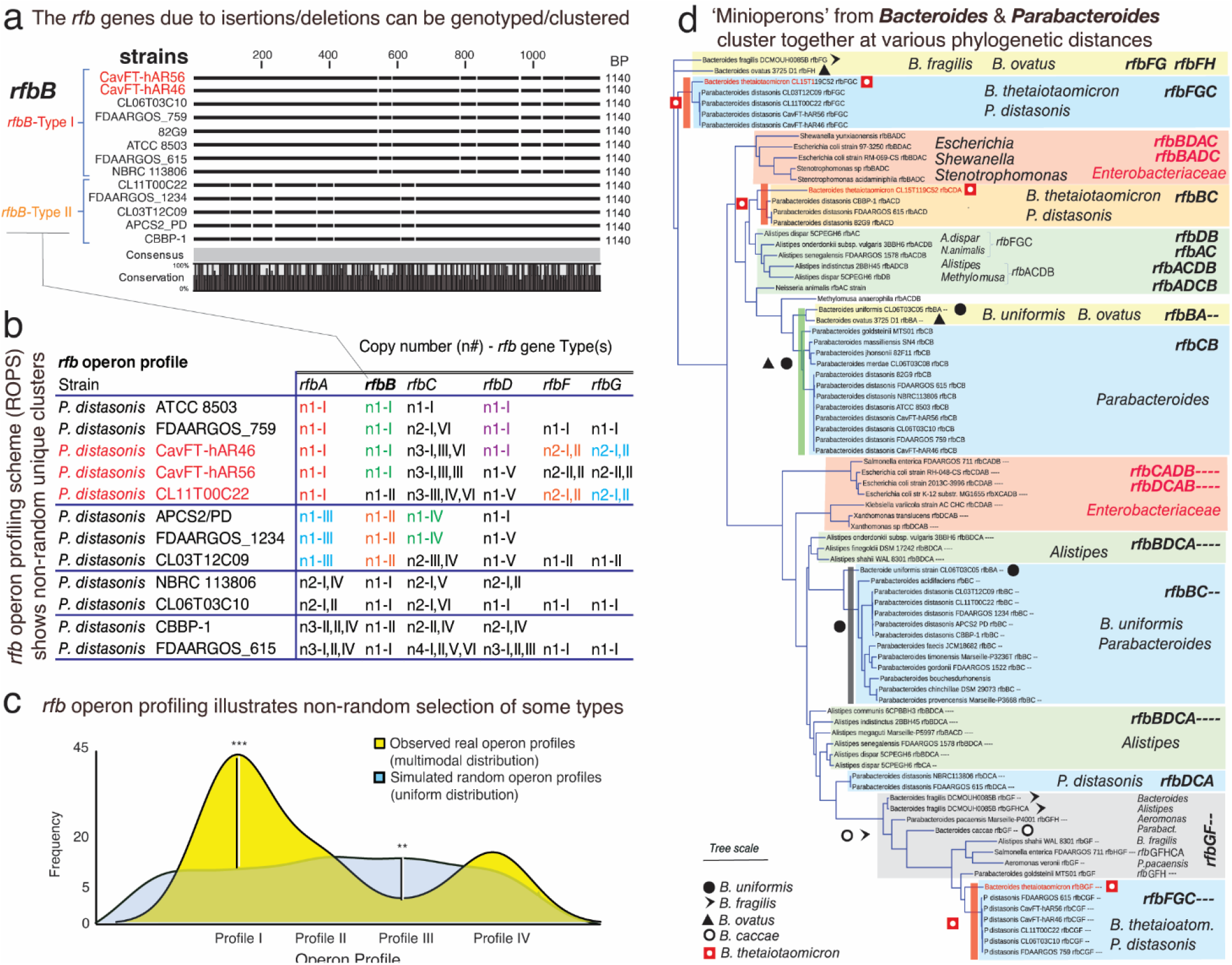
‘*Rfb*-Operon Profiling’ indicates non-random selection and minioperon similarity between *Parabacteroides* and *Bacteroides*, namely *B. thetaiotaomicron*. **A)** Gaps and insertions in *rfb* gene sequences designate different *rfb*-types using protocols described for *rfbA*-typing (3). **Supplementary Figure 4** illustrates the *rfb*-typing of *rfbC/D/F/G*. **B)** Example of global *rfb* operon profiling system (GOPS) for *P. distasonis*. **C)** Density plots between random and real *rfb* operon profiles in *P. distasonis*. Observed types are statistically different from a random (uniform) distribution (**, *** for P<0.05 and P<0.01, respectively). **D)** Phylogenetics across *Bacteroidota* and *Enterobacteriaceae* based on *rfb* mini/operon sequences. Remarkably, several *Bacteroides* species, but namely *B. thetaiotaomicron* CLT5T119C52, cluster together with several *Parabacteroides*, especially *P. distasonis*, irrespective of minioperon considered (red squares; further details in **Supplementary Figures 5**).

### Minioperons from *Bacteroides thetaiotaomicron* and *Parabacteroides* cluster together

Considering the challenges of examining all *Bacteroidota* minioperons with current operon mapping tools (29-32) and the high prevalence of incomplete draft genomes, we next used the NCBI-BLAST database to further investigate if the observed minioperon patterns were *i)* non-random, and *ii)* either widespread across various phyla in the NCBI database or exclusive to particular genera or species. Using *P. distasonis* minioperon sequences, we found that the *rfb* doublets and triplets are limited to *P. distasonis* strains exhibiting ≥99% coverage. Low-matching hits for *rfb* operons/minioperons present with lower (≤24%) sequence coverage were more similar to *Bacteroides* species (*B. thetaiotaomicron, B. caccae, B. cellulosilyticus*) and more distant (≤3% coverage) from *Proteobacteria*, suggesting that the evolution of O-antigen in *Parabacteroides* has been closer to *Bacteroides* than to *Proteobacteria* (**Supplementary Table 2**). Phylogenetic analysis of mini/operons from *Enterobacteriaceae* and *Bacteroidota* with the best BLAST/NCBI sequences (lowest E-scores) further illustrates the well-conserved nature of *Bacteroides* spp. minioperons across genera, and the little similarity to *Proteobacteria* (**Figure 4D**).

Of utmost evolutionary interest, a specific strain of *B. thetaiotaomicron* (CLT5T119C52/USA/c.2018) clustered in three distinct clades (*rfbFGC, BC*, and *FGD*---) with *P. distasonis*, including CavFT strains. This finding indicates that *Parabacteroides/Bacteroides* clusters are likely to have a common ancestor or high affinity for DNA exchange and horizontal ‘operon’ transfer (**Figure 4D**). These *Parabacteroides*/*Bacteroides* minioperon clades also harbored *B. uniformis* or *B. ovatus*, but not *Alistipes* or *Enterobacteriaceae*.

### Operon fragmentation through ‘foreign DNA’ insertions from *Bacteroides*

Given the known high-frequency of horizontal gene transfer (HGT) between *Bacteroides* and gram-negative microbes in the gut (33), the presence of *rfb* minioperons in *Bacteroidota* could indicate that sharing certain *rfb* patterns could reflect adaptation/selection benefits. However, such HGT exchange theory could not explain the observed genus-specificity and low frequency of minioperons in other taxa. Therefore, we hypothesized that the cumulative insertion of ‘foreign DNA’ between *rfb* genes could account for the fragmentation of operons and the variable *rfbFGC->rfbA* separation distances observed in **Figure 2A-D**. We further hypothesized that the genetic exchange in *Bacteroidota* could be a specialized event, confined to niches with compatible species, rather than occurring randomly in the gut. Supported by clades in **Figure 4D** (containing *Bacteroides* matching *Parabacteroides* of CavFT origin), this exchange could explain the low representation of *rfb* minioperons in NCBI. Testing these hypotheses, we first performed whole genome rearrangement analysis to quantify the percentage of genome matching sequences and their fragment sizes. We then examined the sequences located within fragmented *rfbFGC->rfbA* minioperons and select surface antigenic fimbriae inter-minioperon regions as depicted in **Figure 5A**.

**Figure 5.**
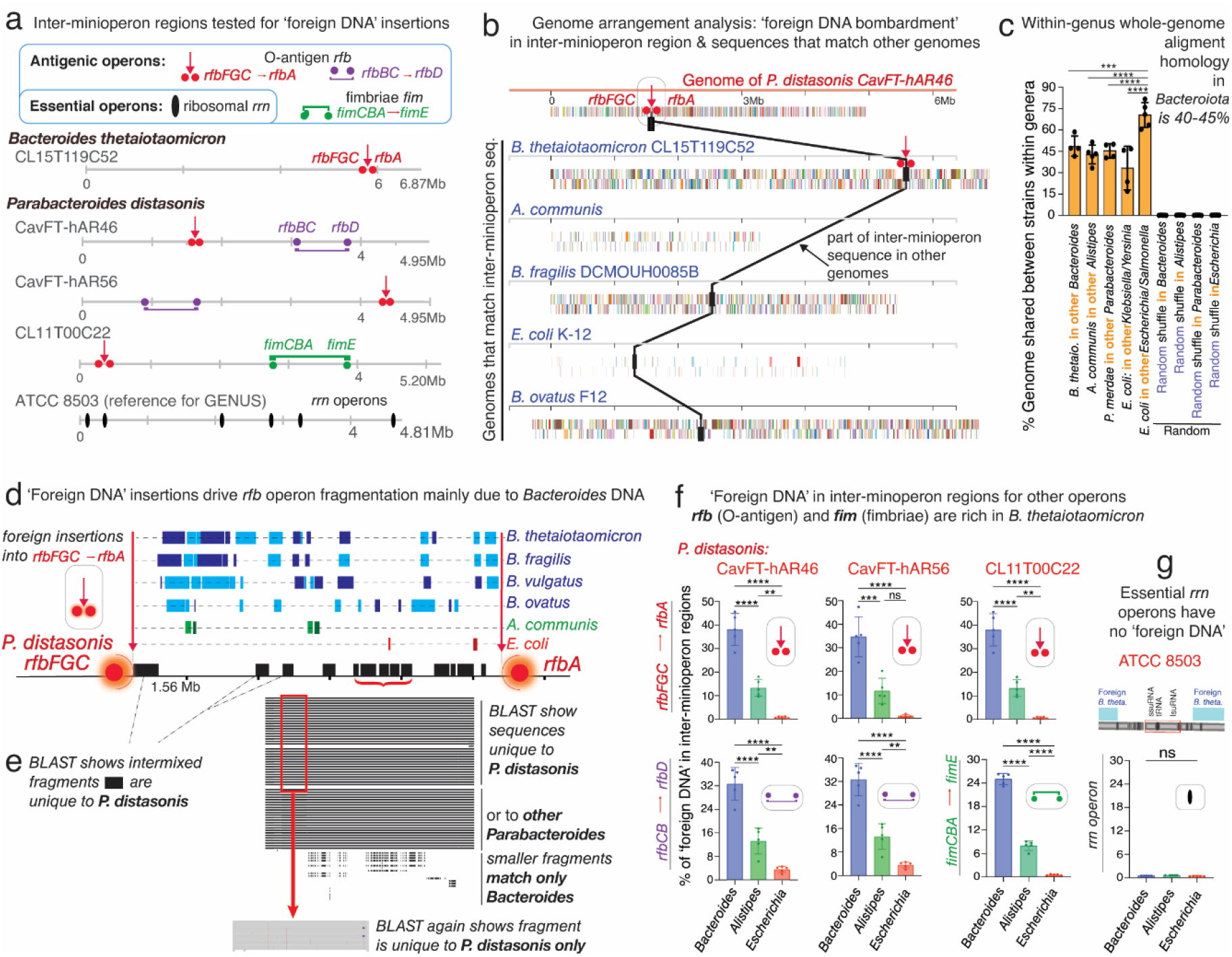
Fragmentation of nonessential operons is driven by insertions of ‘foreign DNA’, mainly *Bacteroides*. **A)** Schematics of O-antigen (*rfb*), fimbriae (*fim*) and ribosomal (*rrn*) gene operons tested for DNA insertions. **B)** Example of *P. distasonis* inter-minioperon DNA fragment found in other genomes. **C)** % of DNA from one species common to the genome of 4-5 other species within the genus. **D)** Schematics showing source of ‘foreign DNA’ insertions into the *rfbFGC*->*rfbA* space separating the *rfb* genes in *P. distasonis*. **E)** ‘Pure’ *P. distasonis* DNA fragments intermixed within inter-minioperon ‘foreign DNA’ insertions. **F)** Genus sources and % of ‘foreign DNA’ fragmenting the *rfb* and *fim* minioperons in *P. distasonis*, as in **Figure 5A**. *Bacteroides* (*B. thetaiotaomicron, B. fragilis, B. ovatus, B. vulgatus*) are the main source of ‘foreign DNA bombardment’ (P<0.0001 vs. *Alistipes* and *Escherichia*). Additional information is available in **Supplementary Table 3. G)** Ribosomal operons (*rrn*, deemed essential) are not fragmented in *P. distasonis*, despite presence of ‘foreign DNA’ in vicinity. *, **, ***, for P<0.01, P<0.001, P<0.0001, respectively. **Supplementary Table 4** shows *rrn* operons are not fragmented in other genera, *i*.*e*., *Prevotella, Bacteroides, Alistipes*, and *Porphyromonas*.

Genome rearrangement analyses showed that 30-45% of the *P. distasonis* genome sequence, in various fragment sizes, match that of *B. thetaiotaomicron*, confirming the two strains belong to different clades at genome scales. However, such finding also emphasizes the potential for disruptive random ‘foreign DNA’ insertions into operons, since genome alignment showed that the largest shared DNA fragment was also present in other *Bacteroides* (**Figure 5B-C**). Of mechanistic interest, we found that the DNA present in the inter-minioperon *rfbFGC->rfbA* sequences represent an overlapping mixture of DNA that matched primarily *Bacteroides* (27-45%; *B. thetaiotaomicron, B. fragilis, P. ovatus*), with limited similarity to *Alistipes* or *E. coli*, and no evidence of similarity to random sequences (shuffled genomes; **Figure 5C-D**). Notably for *P. distasonis*, the entire inter-minioperon sequence represents the combination of ‘foreign’ *Bacteroides* DNA intermixed with DNA sequences verified by BLAST as pure *P. distasonis* (**Figure 5D-E**). This suggests that, over time, such inter-minioperon sequences have either become specific for *P. distasonis*, or they represent insertions of ‘foreign DNA’ from other *P. distasonis*. Examination of *rfb* inter-minioperon regions in other strains and in the antigenic fimbriae (*fim*) minioperons in *P. distasonis* strain CL11T00C22 (strain with most complete set of *fim* genes (34), **Figure 5F**) confirmed the same pattern of *Bacteroides* predominance.

### No fragmentation in ribosomal operons

Lastly, to determine if operon fragmentation also affected essential genes, we examined ribosomal *rrn* operons (∼5000bp containing highly conserved 16S, 23S and 5S rRNA/tRNA genes). We *i)* assessed *rrn* operon integrity, and *ii)* quantified the amount of *B. thetaiotaomicron* DNA found in or near the *rrn* operons of diverse genomes; as an analytical control, we also quantified the amount of *B. thetaiotaomicron* DNA found in randomly generated 5000bp-loci. Analysis revealed that none of the *P. distasonis rrn* operons studied (encompassing 21 rRNAs and interspersed tRNA genes) were fragmented or contained *B. thetaiotaomicron* (Fisher’s exact P=0.0081, vs. randomly selected loci; 0/7 vs. 12/20; **Figure 5G**), despite being exposed to the same ‘foreign DNA insertion pressure’, which was inferred by comparing the average distance of the nearest *B. thetaiotaomicron* DNA insertion to the *rrn* operon (vs. distances to randomly generated locus coordinates; T-test P=0.79). Interestingly, multiple large spans (50,000+bp) of bacterial genome were found to be free of *B. thetaiotaomicron* DNA which could be considered as a future strategy to identify essential operons.

### ‘Foreign DNA’ insertions in *Bacteroidota* on metagenomic statistics

Ranking analysis of all DNA fragments from 15 genomes that aligned to *P. distasonis* CL11T00C22, shown as dot plots in **Figure 6A**, illustrates that *Bacteroides* are the most likely sources of recent genomic exchange with *Parabacteroides* based on the number and size of homologous DNA fragments. Especially intriguing is that *B. thetaiotaomicron* strain CLT5T119C52, again, has the most distinctive sharing pattern (largest number/longest fragments) with *Parabacteroides*, suggesting that DNA exchange is an active, still ongoing interspecies process. Similar insertions of *B. thetaiotaomicron* DNA were also observed in other inter-minioperon regions in other genera (*B. fragilis, Barnesiella viscericola, P. intermedia*, and *A. shahii*). Analysis indicates that not all species are equally likely to receive *B. thetaiotaomicron*, further supporting that inter-species exchange is not random across the phylum, but rather, driven by yet unknown pairing factors among species that share environments where other *Bacteroides* may be DNA donors (*B. fragilis, B. ovatus*; **Supplementary Figure 6 & Supplementary Table 5**).

**Figure 6.**
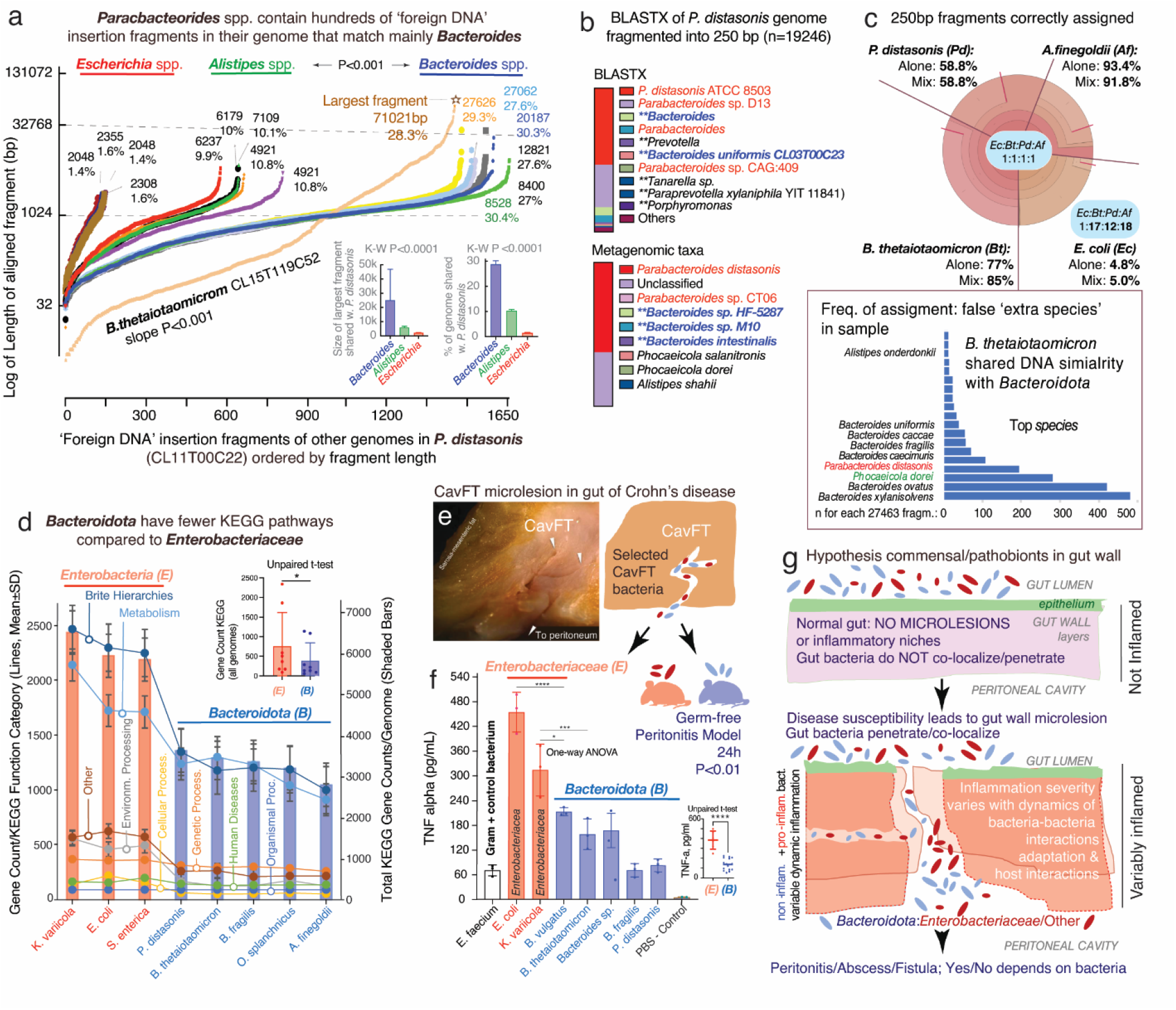
Impact of ‘Foreign DNA’ insertions in *Bacteroidota* on metagenomics, antigenic operons and bacteria-bacteria inflammatory interactions in a model of gut microlesions. **A)** Plot of DNA fragments that align between *Parabacteroides* CL11T00C22 and *Bacteroides, Alistipes, Escherichia* genomes (n=16) ordered by length of each aligning fragment. Notice genus-genus differences. *B. thetaiotaomicron* CL15T119C52 has unique pattern of abundant fragments (maximum fragment sizes and average % of genome shared, inset bar plots). **B)** BlastX (protein) and metagenomic (nucleotide) taxonomic analyses of 250bp-fragmented *P. distasonis* genome. Notice ‘extra species’ assigned by metagenomics (inflation), reflecting ‘foreign DNA’ insertions/exchange across *Bacteroidota*, and not real presence of species (inset bar plot). The n of ‘extra species’ varied with fragment length (Pearson corr. 0.84, P<0.05). The performance of BLAST and BLASTX depends on bacterial genome (**Supplementary Figure 8**). **C)** Metagenomic community simulation with *Bacteroides, Parabacteroides, Alistipes*, and *Escherichia* (1:1:1:1 genomes). Krona plot (relative abundances within hierarchies of metagenomic classifications (35)) illustrates *E. coli* sequences are poorly assigned to *E. coli* leading to relative ratio overestimation of *Bacteroidota* abundance (1:17:12:18; see Krona plots for individual genomes (‘Alone’) in **Supplementary Figure 9**). *Bacteroides* is commonly listed as ‘extra species’ (bar plot; complete list in **Supplementary Figure 7**) **D)** KEGG pathway and total gene counts in *Enterobacteriaceae* and *Bacteroidota*, highlighting the significant differences for *Bacteroidota* (details in **Supplementary Table 7**). **E)** Glycostaining of LPS extracted from *E. coli* and *P. distasonis* cultured in different media. **F)** Average pro-inflammatory cytokine secretion by bacteria in the stimulation, indicating that *Bacteroidota* release less pro-inflammatory cytokines compared to *Enterobacteriaceae*. **G)** Hypothetical model of gut microlesions with colonization of commensal/pathobionts modulating inflammation. Peritonitis model showed mice with *Enterobacteriaceae* had fatal peritonitis, but not if receiving *Bacteroidota B. thetaiotaomicron, B. fragilis*, or *P. distasonis*. *P<0.01; ****P<0.00001.

With a large number of ‘foreign DNA’ insertions in the genome that could disrupt operons (n=1450-1650, x-axis **Figure 6A**), there is also potential for impacting metagenomic results. In examining the taxonomic assignment of inter-minioperon sequences using BLAST, we first illustrated the potential for metagenomic overestimation (‘inflation’: calling of ‘extra species’ in a sample when they are not there). BLAST suggests this could be important for strains avid to share DNA, but not for non-avid strains. While numerous inter-minioperon sequences matched to several *B. thetaiotaomicron* strains in NCBI with 100% coverage and >99% identity (81% of top hits), in a few cases (<4%) DNA matched other *Bacteroides* with lower similarity (<85%), such as *B. longzhouii* or *B. faecis*, and *B. fragilis* (21.4% top hits, **Supplementary Table 6**). Together, as illustrated in **Figure 5D-E**, analyses indicate DNA insertion similarity is more likely with *Bacteroide*s, being restricted to *Bacteroidota*. To expand these BLAST-derived inferences, we used the BV-BRC metagenomics workflow for taxonomic classification of DNA sequences ‘in bulk’ to inspect entire genomes. By fragmenting the genome of selected strains into equal nonoverlapping 250bp simulated ‘reads’, we determined, *in-silico*, that the species inferred from inter-minioperon sequence queries in NCBI were also reproducible by metagenomics.

Since metagenomics uses sequences to deduce i) taxonomic composition using BLAST (nucleotide database), or ii) taxonomic composition and pathways/functions using BLASTX (protein database), ‘foreign DNA’ insertions in *Bacteroidota* affect the accuracy of these tools. The impact of ‘foreign DNA’ insertions can be visualized for both ‘individual absolute metagenomic’ analyses (fragmented genomes analyzed individually, **Figure 6B**), and on ‘relative metagenomic analyses’ for a simulated community with four genomes (*A. finegoldii, B. thetaiotaomicron, P. distasonis*, and *E. coli*, 1:1:1:1, 1X and 20X, **Figure 6C**). Although metagenomics aligns fragments/reads into contigs (algorithm assumption) and then finds their best match in a database with genes representing selected reference strains (BV-BRC n=24868; 16840 species; 30573 taxon units), analysis shows the presence of DNA from numerous *Bacteroidota*, especially the species shown by NCBI-BLAST in inter-minioperon regions. Top species for *B. thetaiotaomicron* included *B. xylannisolvens, B. ovatus, P. dorei* and *P. distasonis*, further confirming inter-species affinity for selective genetic sharing. Diagnostically relevant, and as anticipated, results also illustrate that metagenomic workflows could overestimate diversity by suggesting the presence of ‘extra species’ not actually present in a sample. Of note, the calling of ‘extra species’ by metagenomics depends largely on the fragment length used; however, while the number of fragments called ‘extra species’ is reduced as fragments get longer (from 100bp to 16000bp), the relative presence of ‘extra species’ increases with fragment length (Pearson’s P<0.05), confirming overestimation and suggesting the need of algorithm revision in current workflows.

Assessing communities in confined micro-niches will remain challenging using current metagenomics, since underestimation of present species (‘deflation’: species underestimated in abundance or deemed absent from sample) could also occur in cases where pathobionts/commensals coexist. For example, our relative community analysis showed that if a species is largely under-assigned by metagenomics (*E. coli*, ∼4%, **Figure 6C**), there will be further overestimation of *Bacteroidota* ratios (12-to-18-fold magnification vs. *E. coli*, **Figure 6C** and **Supplementary Figure 7**).

### Reduced KEGG pathways, TNF-alpha induction, and peritonitis by (CavFT) *Bacteroidota*

Since a widespread process of DNA insertions throughout the genome could affect the functionality of various operons and genetic pathways, we next tested if such a phenomenon of operon disruption could be visualized by observing a lower number of functional pathways using a pathways database. By using the Kyoto Encyclopedia of Genes and Genomes (KEGG), a large-scale molecular dataset generated by genome sequencing, high-throughput experiments, and manual curation to infer enzymatic pathways across bacterial genomes (36), we confirmed that *Bacteroidota* have significantly less pathways than *Enterobacteriaceae* (T-test, P<0.05). Although the database may be biased towards *Enterobacteriaceae* due to more published evidence, **Figure 6D** demonstrates that selected *Bacteroidota* have significantly fewer functional pathways responsible for ‘metabolism’ and ‘environmental information processing’, while there was no difference for pathways responsible for ‘organismal systems,’ ‘human diseases,’ ‘genetic information processing,’ and ‘cellular processes’, supporting that the findings are well-controlled for basic pathways functions.

Of interest, when testing representative CavFT bacterial isolates derived from the gut wall of patients with CD in our laboratory (**Figure 6E**), we determined that the overall inflammatory potential of bacteria on murine macrophages was significantly reduced compared to *Enterobacteriaceae*. **Figure 6F** shows that TNF-alpha production by macrophages cultured *in vitro* (RAW264.7 cells) exposed to heat-treated bacterial extracts was about half the immune-proinflammatory potential observed for *E. coli* and *Klebsiella variicola*. Controlling for the apoptotic effect that bacterial extracts could have on macrophages at different extract dilutions, (measured using cell viability MTT assay), TNF-alpha data shows that *Bacteroidota* isolates from CavFT have non-inflammatory antigenic phenotypes compared to *Enterobacteriaceae* (*e*.*g*., O-antigen *rfb* operons) or *Enterococcus faecium* (gram-positive control). To further validate *in vivo* the sub-inflammatory potential of CavFT *Bacteroidota*, we injected suspensions of live bacteria into the peritoneal cavity of germ-free Swiss Webster mice to quantify their inflammatory potential. Our peritonitis model, based on 270,000 real-time telemetry data points, revealed that the mice receiving *E. coli* or *K. variicola* became febrile, then hypothermic, lethargic, and moribund within 24h post injection, while mice receiving CavFT *Bacteroidota* only became transiently hypothermic following the injection, overall being clinically normal or telemetrically less active until the end of study 24h post injection.

Based on evidence of antigenic operon fragmentation by *Bacteroides* and the reduced proinflammatory potential of *Bacteroidota in vitro and in vivo*, **Figure 6G** depicts a hypothesis where *Bacteroidota* invade, interact, and adapt to gut wall micro-niches where other enteric bacteria (*e*.*g*., *Enterobacteriaceae* and *Bacteroidota*) may be present and dynamically fluctuate to explain the cyclical remission-flare dynamics of gut wall inflammation and complications in chronic bowel diseases like CD.

## Discussion

To explore the causes of pro-inflammatory surface antigen variability affecting gut pathobiont dualism within *Bacteroidota*, this study initially assessed the integrity of the *rfb* operon utilizing a selected set of complete genomes. Validation of findings was achieved by extending the analysis to other genomes and annotated sequence repositories, mainly using NCBI/BLAST. Of note, *Bacteroidota* have their *rfb* operons either intact (*Odoribacter, Porphyromonas gingivalis)*, duplicated (*Alistipes*) or, mainly, completely or partially fragmented into ‘minioperons’, which can be classified into at least 5 categories using a broadly-applicable operon cataloguing system (**Figure 1C**). Overall, minioperons are highly-conserved features within genera, non-random, rare across NCBI databases, and form distinctive patterns sporadically shared among distant species, indicating a common mechanism for operon fragmentation within the phylum. By characterizing and cataloging *rfb* operon fragmentation patterns, we determined that the insertion of ‘foreign DNA’ from other bacteria, mainly *Bacteroides*, could explain operon fragmentation as observed in *P. distasonis* (supernumerary minioperon fragmentation), which was used together with *Alistipes* spp. as model bacteria for hypothesis testing.

Intriguingly, the presence of *Bacteroides* ‘foreign DNA’ insertions between antigenic operon genes (*rfb, fim*), but not essential ribosomal operons (*rrn*), implies that DNA insertions/operon damage favors selection if effects allow bacterial survival. Indeed, previously observed patterns of operon disruption in *Bacteroides* spp. have been linked to functional themes related to niche-habitat survival (37, 38), suggesting a similar phenomenon may occur across *rfb* operons in *Bacteroidota*. Furthermore, prior literature supports that, in addition to reductions/loss of the O-antigen (39, 40), *rfb* gene mutations influence bacterial survival (41) and bacteriophage infection (42). While more research is needed to elucidate how *rfb* gene dosage and structure influence LPS and/or other cellular KEGG ontology maps and functions in *Bacteroidota*, our analyses demonstrated the lack of O-antigen production by *P. distasonis* under different growth conditions and the lower TNF-alpha production induced by CavFT *Bacteroidota* isolated from CD patients in our laboratory (14, 15), or the lack of induction of peritonitis, which contrast reports of severe peritonitis due to *B. thetaiotaomicron or B. fragilis*, which are commonly seen in immunocompromised individuals (43-46).

Of evolutionary interest, several *Bacteroides*, including the bacterium *B. thetaiotaomicron* CL15T119C52 (human/feces/2018) were remarkably noted to cluster with *P. distasonis* CavFT strains (CD patients/2019) based on conserved minioperons (**Figure 4**). This suggests there is preferential DNA exchange among certain strains (namely *Bacteroides* as shown throughout the study, **Figures 5 and 6A**). Our *in-silico* analyses, manual annotation, and experimental observations serve as a proof-of-principle for the (primarily) *Bacteroides* DNA insertion mechanism of operon fragmentation and its potential impact on metagenomics. Findings provide novel insights and opportunities for diagnostics and therapeutic developments, especially considering that the most remarkable findings, such as the distancing of the *rfbFGC*->*rfbA* minioperons, involve bacteria isolated from chronic inflammatory microenvironments in CD patients. Our study for the first time examines and reports the genomic features of CavFTs in context with other members of the phylum, also isolated from CavFTs, which could evolve and adapt into lineages that may survive on/inside micro-niches in the inflamed gut wall (23).

In conclusion, our findings highlight that operon fragmentation provides novel mechanistic insights for commensal adaptation and metagenomic applications. An improved understanding of *Bacteroidota rfb* operons can provide valuable insights into bacterial genetics and their role in human health, as well as help refine experimental strategies for studying host-pathogen interactions. Future studies on these interactions or disease causality and operon integrity would benefit from combining bacterial isolation with genomic and transcriptomic sequencing to assess KEGG-pathways functionality.

## Materials and methods

### Genome Databases and Data Collection

Genome wide genetic analyses for the *rfb* operon were conducted on publicly available datasets and on reference strains sequenced in our laboratory that we isolated from intramural cavernous lesions in the damaged bowel of Crohn’s disease patients as previously reported (5, 23). Only complete reference genomes of selected strains of human derived *Enterobacteriaceae (e*.*g*., *E. coli* K12, as reference for the *rfb* operon), selected strains for representative genera of the *Bacteroidota* phylum, all available reference strains for all species within the *Alistipes* and *Parabacteroides*, and all available strains for the *Parabacteroides distasonis* species were used in this study. National Center for Biotechnology Information (NCBI) GenBank and the Bacterial and Viral Bioinformatics Resource Center (BV-BRC) were accessed for bacterial genomic data. Data collected from each genome includes accession number, genome length, *rfb* gene copy number, nucleotide sequence, location, and direction within the genome. All results of the *rfb* gene query were manually verified using NCBI annotated graphical portals. Additional data was collected to find the closest homolog of conserved minioperons in *P. distasonis*; We used the previously obtained FASTA sequences of *P. distasonis rfb* minioperons as input queries in the nucleotide Basic Local Alignment Search Tool (BLAST) using default settings under the nr/nt database, and BLAST hit result data and *rfb gene* sequence data for matching organisms from homologue queries were subsequently collected.

### Visualization of *rfb* gene dosage and location

The *rfb* genes for each genome were graphed (using gene coordinates) along a linear axis representing each bacterial genome to visualize the respective *rfb* operon distribution pattern in each bacterium. *Rfb* gene and minioperon/operon copy numbers for each genome were used to generate heatmaps demonstrating relative *rfb* gene dosages. Heatmaps for *rfb* gene dosages in *P. distasonis* strains were created using ClustVis (47) and a summary heatmap of *rfb* gene dosages for all genomes was generated with the web tool Morpheus (https://software.broadinstitute.org/morpheus).

### Homology Analyses

Sequences of recurring doublets; *rfbCB*(++) and *rfbBC*(--), and of recurring triplets; *rfbACD*(+++), *rfbDCA*(---), *rfbFGC*(+++), and *rfbCGF*(---), were collected and aligned with both their respective doublets/triplets in *Parabacteroides* genomes as well as their respective reverse complements. In *Alistipes* spp., sequences of its *rfbACDB*(++++)/*BDCA*(----) operons were aligned to determine homologies at the species level. Of note, no consistent patterns of minioperons were observed in the selected *Bacteroides* or *Prevotella* spp., thus no analyses could be performed to determine their respective homologies. Sequence homology was performed using the Sequence Identity and Similarity (SIAS) tool (http://imed.med.ucm.es/Tools/sias.html).

### Global *rfb* operon profiling system (GOPS) design

DNA sequences for each *rfb* gene in complete *P. distasonis* strain genomes were collected and aligned respectively (*e*.*g*., *rfbB* gene alignment, *rfbC* gene alignment, etc.) in CLC Genomic Workbench (commercially available). Following our previously developed *rfbA*-typing protocol (3), *rfb*-gene-types were designated for each *rfb* gene alignment. The aggregate results of the copy number(s) and *rfb*-type(s) for each genome were used to construct an example *rfb* operon profiling system, utilizing the nomenclature system previously proposed for *rfbA*-type reporting in *Bacteroidota* (3).

### Intergene and *rfb* loci statistics

The linear distance between each *rfb* gene was determined by calculating the number of base pairs in between consecutive genes in each genome. *Rfb* loci were counted in each genome, where a locus was determined to be a discrete location of either a single *rfb* gene or a cluster of contiguous *rfb* genes (operons/minioperons). Statistical analyses were performed to determine if *rfb* intergene linear distances and number of *rfb* gene loci differed significantly between the genera and phyla examined. The range and variance of *rfb* intergene distances was also determined at the genus and species level for *Parabacteroides* and *Alistipes*. Prior to statistical analysis, *rfb* intergene linear distance data was log transformed. Transformed *rfb* intergene linear distances and *rfb* gene loci data were analyzed using Brown-Forsythe ANOVA and Welch’s ANOVA to determine if statistically significant differences existed between the respective mean values of these data for *Enterobacteriaceae, Parabacteroides, Bacteroides, Prevotella*, and *Alistipes*.

### Construction of phylogenetic trees

The phylogenetic tree of whole genomes was made by Bacterial and Viral Bioinformatics Resource Center (BV-BRC), under the default setting for codon tree (which uses 100 amino acids) and nucleotide sequences from BV-BRC’s global Protein Families to build an alignment and then generate a tree based on the differences within those selected sequences. For the phylogeny of *rfb* gene clusters (operons and minioperons), the nucleotide FASTA sequences encoding the *rfb* operons and minioperons were downloaded from NCBI database. The multiple sequence alignments of all nucleotide sequences were performed using the clustal omega (https://www.ebi.ac.uk/Tools/msa/clustalo/) which was used for the construction of phylogenetic trees with the maximum likelihood methods for evolutionary analysis by using Webserver IQ-Tree (http://iqtree.cibiv.univie.ac.at/) under default parameters of ultrafast bootstrap. The phylogenetic trees with branches were built with iTOL (https://itol.embl.de/).

### Primer design for amplification of the *Alistipes* spp. *rfbBDCA/ACDB* operon

Primer design was conducted by identifying left and right flanking regions of the *Alistipes* spp. *rfbBDCA/ACDB* operon alignment of which were whole (*i*.*e*., no gaps or deletions) throughout all sequences. Then, from the corresponding regions of the *rfbBDCA/ACDB* operon sequence alignment consensus sequence, left and right flanks of approximately 20 base pair sequences were selected and entered into the Basic Local Alignment Search Tool (BLAST) to confirm accuracy in identifying *Alistipes* spp. utilized in this study.

### Random sequences

Random generation of genomes and the random shuffling of complete genomes were conducted using Sequence Manipulation Suite software (https://www.bioinformatics.org/sms2/shuffle_dna.html) with parameters that matched the %GC content of relevant and selected species, including 43% GC content to mimic *Bacteroides* genomes (*B. thetaiotaomicron* CL15T119C52: 43.07%, *B. fragilis* DCMOUH0085B: 43.61%, *B. ovatus* F-12: 41.98%, *B. vulgatus* NCTC10583: 42.03%, and *B. dorei* MGYG-HGUT-02478: 42.04% for an average of 42.55%).

### Calculation of the gene combination of *rfb* minioperons

For the bacterial genomes used in this study, the number of potential *rfb* minioperon combinations for each bacterium was determined by calculating the sum of the permutations of these minioperon gene combinations.

### LPS Extraction and Glycoprotein Staining

*P. distasonis* and *E. coli* were cultured in five different conditions (BHI+ 5% Yeast Extract, 50% BHI Broth, BHI Broth+5% Yeast Extract with normal person heat-killed feces, BHI+ 5% Yeast Extract with 4% bile and 50% BHI Broth with CD patient heat-killed feces) and centrifuged at 2000g for 10 minutes. The supernatant was decanted and the masses of the pellets were obtained. The LPS was extracted by LPS extraction kit (Sigma Aldrich Catalogue Number: MAK339) using the manufacturer’s instructions. In short, the cell pellets were resuspended in lysis buffer at a ratio of 100 μL of buffer for every 10 mg of cell pellet. The bacterial cells were lysed using a MP Fast-Prep 24 Homogenizer, using 4 rounds of shaking at 4 m/s for a duration of 20 seconds. The lysed pellets were then centrifuged at 10000g for 5 minutes to sediment the cell debris. The supernatant containing LPS was collected in a separate tube, to which proteinase K was added for a final concentration of 0.01 mg proteinase K/ml. The solution was heated to 60°C and held for 60 minutes. Following this, the solution was centrifuged at 10000g for 5 minutes, and the supernatant containing the free LPS was collected. The LPS extracts were loaded into a NuPAGE 4-12% Bis-Tris Gel (NP0335BOX) along with commercially available LPS (Sigma) derived from *E. coli*, as well as a pre-stained protein ladder. The gel bands were fixed by incubating the gel in 50% Methanol for 30 minutes at room temperature. The gel was stained using a commercially available Pierce™ Glycoprotein Staining Kit according to the manufacturer’s instructions.

Briefly, the gel was then immersed in a 3% acetic acid solution and incubated for 10 minutes at room temperature. The solution was removed and replaced with fresh acetic acid for another 10 minutes. The gel was then submerged in the oxidizing solution and incubated for 15 minutes with agitation using an orbital shaker. The gel was washed three times with 3% acetic acid for minutes. The gel was then transferred to the glycoprotein staining solution and incubated for 15 minutes on an orbital shaker.

### TNF-α Stimulation Assay in RAW 264.7 cells

The RAW 264.7 cells were flushed and cultured in DMEM (Dulbecco’s Modified Eagle’s Medium (DMEM)) and supplemented with 10% FCS, NEAA, glutamax, penicillin–streptomycin. Cells were plated for experiments after 6 days. Cells were plated at 4 × 10^4^ cells per well of a 96-well plate, and all cell lines were seeded 16 hours prior to challenging. The bacteria were cultured, and pellets were resuspended in PBS. Each pellet was heat killed by placing in a heating block for 30 min at 95°C. To normalize the concentrations of each bacterial suspension, the OD600 value was taken. The different dilutions (1:1, 1:5, 1:25 and 1:125 dilution) of the heat killed bacterial extract were added to the RAW 264.7 cells, then the medium was collected after 18 h for testing by TNF enzyme-linked immunosorbent assay.

### Peritonitis model

To quantify the impact of bacteria on the ability to trigger inflammation *in vivo*, we conducted studies with mice using a peritonitis model. Using fresh anaerobic bacterial preparations (10^8 CFU/mL), each animal received an intraperitoneal injection of selected bacteria and underwent continuous monitoring for 24h prior to being euthanized. Real time mobility in the cage measured with subcutaneous RFID tags, response to stimuli, body temperature, mortality, and bacterial viability in the peritoneal fluid were measured as main outcomes.

### Statistics

Data analysis was conducted using parametric and non-parametric statistics using parametric or non-parametric methods (*e*.*g*., student T-tests, ANOVAs) using the software STATA (v17), R, and GraphPad, which was used primarily to make bar plots or boxplot illustrations where significance is represented as asterisks based on the level of significance which was held at p<0.05(*), <0.001(**) and <0.0001(***). Stata functions were used to assess multimodality (minioperon types observed vs. simulated random) as previously reported in our laboratory (48). Together, data analysis represents over 636 analytical submissions made to the various bioinformatic resources described in this study.

### Assessing operon fragmentation

Using the Bacterial and Viral Bioinformatics Resource Center (BV-BRC; Bacterial and Viral Bioinformatics Resource Center) as the data source and genome alignment tool, genomic homologies were first assessed between *P. distasonis* and representative members of *Bacteroides, Alistipes*, and *Escherichia*, respectively, to test the hypothesis that genomic insertions were not from random sources but rather derived primarily from *Bacteroides*. We then identified unique *rfb* order series that were present across seemingly unrelated genomes (chronologically, geographically), including the series ‘*rfbFGC* ⟶ *rfbA’*, wherein a triplet (*rfbFGC*) is followed by a nearby separated singlet gene (*rfbA*). Focused on *Bacteroides* and *P. distasonis* as recipient genomes, including CavFT strains, we examined inter-minioperon DNA sequences (*e*.*g*., between *rfbFGC* and *rfbA)* to conduct genome-wide arrangement analysis between such regions in *P. distasonis* and the genomes of *B. thetaiotaomicron*, other *Bacteroidota*, and *E. coli* to quantify their genus-dependent potential as donors of ‘foreign DNA’. Intra-phylum genome homologies of *Enterobacteriaceae* were also measured as a referent. We assessed the genetic homologies of three inter-minioperon segments (*rfbFGC* ⟶ *rfbA, rfbCB* ⟶*rfbD, fimCBA* ⟶*fimE*) present either completely or uniquely across three *P. distasonis* strains (CavFT-hAR46, CavFT-hAR56, and CL11T00C22). Whole genome homologies were then compared to inter-minioperon genetic homologies.

Next, selectivity of operon fragmentation was assessed in the *P. distasonis* ATCC 8503 reference genome, for which 7 16S rRNA loci were annotated in BV-BRC. These 7 loci were compared with 20 randomly generated loci of similar size to compare the amount of *B. thetaiotaomicron* DNA in or near each locus.

### Genome fragmentation alignment analysis

We quantified and plotted the distribution of DNA fragments detected by genome alignment ranked by fragment number (x-axis) vs. the size of each fragment (y-axis) across 16 genomes (4 *Escherichia* spp., 5 *Alistipes* spp., and 7 *Bacteroides* spp.) individually aligned to *P. distasonis* in BV-BRC.

### Assessment of fragmentation in other antigenic (fimbriae, *fim*) operons

Recently, researchers investigating the antigenic cell surface structure of *P. distasonis* identified fimbriae (*fim*) operon genes in several strains (34), though there was again ample heterogeneity in the type and quantity of these genes amongst the strains investigated, suggesting operon fragmentation could also occur for other functions. For this analysis we used the *P. distasonis* CL11T00C22 strain genome as it contains the most complete set of identified *fim* operon genes.

### Metagenomics of *Bacteroidota*

Whole genome sequences of select *Bacteroidota* (and *Escherichia coli* K-12 as referent) downloaded from NCBI were split into 250bp sequences using FASTA-splitting software (https://www.bioinformatics.org/sms2/split_fasta.html) to approximate the range of *in vivo* DNA samples which are commonly used for shotgun sequencing (49). The sequences for each respective genome were then used as inputs for taxonomic classification in BV-BRC as well as for assessment of matching protein sequences using BLASTX.

We validated our hypothesis using principles of genomic and metagenomics data analysis using well-referenced and established methods that are readily accessible over the internet to the scientific community (50-52). We used the Bacteria and Virus Bioinformatics Resources Center (BV-BRC) web server as it integrates multiple analysis steps into single workflows (52), by combining the resources from the former Pathosystems Resource Integration Center (PATRIC), the Virus Pathogen Database and Analysis Resource (ViPR) and the Influenza Research Database (IRD) (50, 51). Analysis tools include pipelines for read assembly, open reading frame prediction, and annotation with BLAST, and GO pathways classifiers. The BV-BRC web server is an easy-to-use web-based interface for processing, annotation, and visualization of genomic and functional metagenomics sequencing data, designed to facilitate the analysis of data by non-bioinformaticians (52). Metagenomics results derived from Kraken2, are visualized via Krona as a dynamic online feature for hierarchical data and prediction confidence (35). The BV-BRC online setting provides scientists a fast open-source strategy to analyze raw sequences and generate complex comparative analysis by selecting reference genomes or genomes uploaded by the user to an academic account. Genome arrangement analysis was conducted using the tool ‘Comparative Analysis tools’ available in beta version. Sequence and genome coordinate data can be examined as summary tables or visually through colored interactive arrangement plots. BV-BRC facilitates effortless inspection of gene function, clustering, and distribution. The webserver is available at https://www.bv-brc.org/.

## Acknowledgements

This project was supported by 1) NIH grant R21 DK118373 to A.R.-P., entitled “Identification of pathogenic bacteria in Crohn’s disease.” and 2) Crohn’s & Colitis Foundation Student Research Fellowship Award #932611 to Nicholas Bank, entitled “Molecular Immunology of *Parabacteroides distasonis* in Crohn’s Disease and Role of Strain Diversity in Macrophage Activation. Partial support earlier originated for A.R.-P. from a career development award and recently from a Litwin Pioneers award from the Crohn’s and Colitis Foundation.

