## Supplementary Materials for "The basis of antigenic operon fragmentation in *Bacteroidota* and commensalism"

### Supplementary Figures 1-9

### Supplementary Tables 1-7

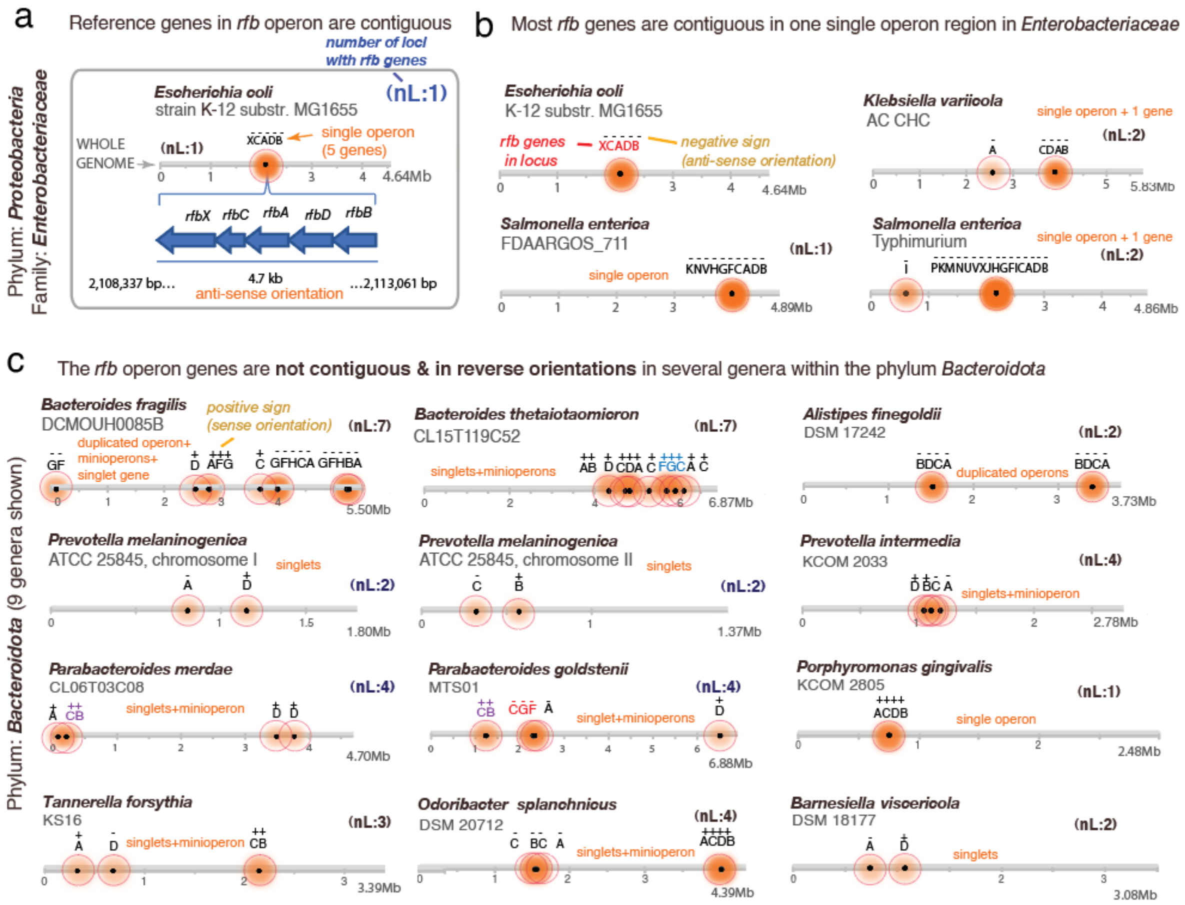

**Supplementary Figure 1. The *rfb* operon in *Enterobacteriaceae* is contiguous, but in *Bacteroidota* it is fragmented into 'minioperons' with diverse orientations or duplications.** Lines represent complete genome with distribution of *rfb* gene clusters/operons (shaded circles) with genomic orientations (+, sense; -, antisense). Circle shading: the darker, the more genes in cluster. nL, number of *rfb* gene loci (clusters or gene singlets) in each genome. **A)** Classical *rfb* operon genes clustered contiguously in reference genome *E. coli* K-12**. B)** The *rfb* operons are single-copied in *Escherichia coli* and *Enterobacteriaceae*. **C)** The *rfb* operon in most genera within the *Bacteroidota* phylum is often fragmented into 'minioperons'. Available complete genomes representing 8 of the 9 genera within *Bacteroidota* illustrate various types of operon fragmentation in the phylum (*Alistipes, Bacteroides, Parabacteroides, Prevotella, Paraprevotella, Barnesiella*, *Tannerella*, *Odoribacter* and *Porphyromonas*).

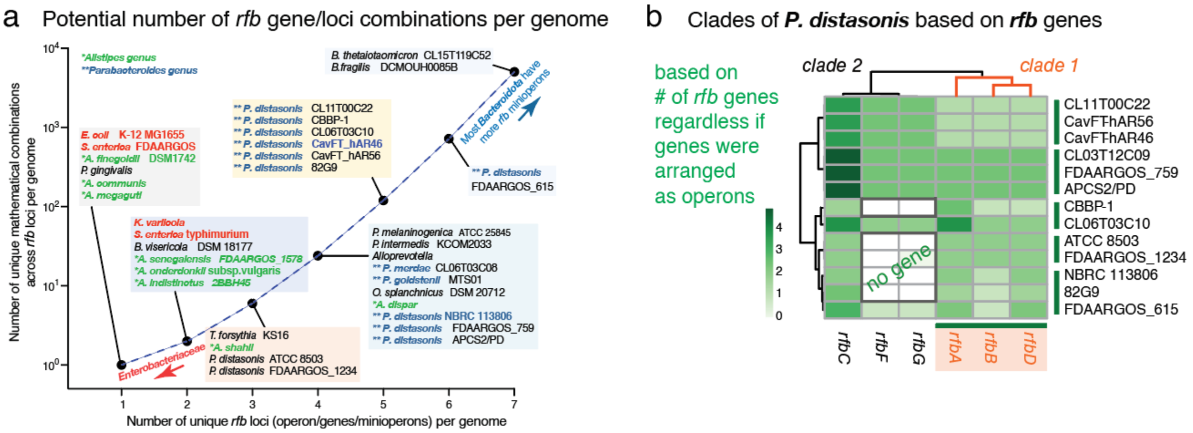

**Supplementary Figure 2. A)** Number of possible *rfb* operon gene combinations based on the unique *rfb* minioperons present in different bacteria used in this study. **B)** Heatmap clustering of *P. distasonis* strains based on *rfb* gene dosage alone, regardless of the presence of minioperon formation.

**a** Pairwise nucleotide sequence similarity across *rfb* minioperons (sense vs. antisense) within the ***Parabacteroides*** genus

Doublet minioperons (column groups: ++ CB: *P. chartae* to *P. goldsteinii*; - - BC: *P. bouchesdurhonensis* to *P. faecis*)

| | | *P. chartae* | *P. distasonis* | *P. merdae* | *P. johnsonii* | *P. massiliensis* | *P. goldsteinii* | *P. bouchesdurhonensis* | *P. provencensis* | *P. chinchillae* | *P. gordonii* | *P. timonensis* | *P. faecis* |
|---|---|---|---|---|---|---|---|---|---|---|---|---|---|
| ++ CB | *P. chartae* | 100 | | | | | | | | | | | |
| | *P. distasonis* | 72.06 | 100 | | | | | | | | | | |
| | *P. merdae* | 72.45 | 76.68 | 100 | | | | | | | | | |
| | *P. johnsonii* | 72.56 | 76.62 | 89.12 | 100 | | | | | | | | |
| | *P. massiliensis* | 73.67 | 76.27 | 88.95 | 90.18 | 100 | | | | | | | |
| | *P. goldsteinii* | 73.11 | 75.04 | 76.38 | 76.67 | 77.02 | 100 | | | | | | |
| - - BC | *P. bouchesdurhonensis* | 72.27 | 73.93 | 74.44 | 74.85 | 73.92 | 76.44 | 100 | | | | | |
| | *P. provencensis* | 73.03 | 74.51 | 75.46 | 75.17 | 74.66 | 78.07 | 80.61 | 100 | | | | |
| | *P. chinchillae* | 72.91 | 74.45 | 75.34 | 75.28 | 74.78 | 78.13 | 80.50 | 99.01 | 100 | | | |
| | *P. gordonii* | 73.14 | 76.41 | 77.70 | 78.58 | 79.11 | 85.72 | 74.83 | 76.71 | 76.77 | 100 | | |
| | *P. timonensis* | 72.90 | 74.60 | 76.36 | 77.58 | 77.47 | 85.66 | 76.12 | 75.83 | 76.12 | 88.12 | 100 | |
| | *P. faecis* | 73.29 | 74.98 | 76.61 | 76.73 | 76.73 | 86.76 | 74.98 | 76.38 | 76.73 | 83.96 | 84.43 | 100 |

**b** Pairwise nucleotide sequence similarity across *rfb* minioperons (sense vs. antisense) within the ***Alistipes*** genus

Quadruplet minioperons (column groups: ++++ ACDB: *A. indistinctus* to *A. senegalensis*; - - - - BDCA: *A. indistinctus* to *A. dispar*)

| | | *A. indistinctus* | *A. onderdonkii* | *A. senegalensis* | *A. indistinctus* | *A. finegoldii* | *A. onderdonkii* | *A. shahii* | *A. communis* | *A. senegalensis* | *A. megaguti* | *A. dispar* |
|---|---|---|---|---|---|---|---|---|---|---|---|---|
| ++++ ACDB | *A. indistinctus* | 100 | | | | | | | | | | |
| | *A. onderdonkii* | 64.71 | 100 | | | | | | | | | |
| | *A. senegalensis* | 71.28 | 77.60 | 100 | | | | | | | | |
| - - - - BDCA | *A. indistinctus* | 27.50 | 36.74 | 30.47 | 100 | | | | | | | |
| | *A. finegoldii* | 28.83 | 38.56 | 30.82 | 60.38 | 100 | | | | | | |
| | *A. onderdonkii* | 35.88 | 35.37 | 37.80 | 60.23 | 77.77 | 100 | | | | | |
| | *A. shahii* | 28.64 | 37.34 | 28.65 | 59.13 | 67.69 | 68.46 | 100 | | | | |
| | *A. communis* | 32.68 | 35.10 | 35.03 | 51.17 | 61.34 | 60.51 | 64.50 | 100 | | | |
| | *A. senegalensis* | 27.52 | 35.38 | 27.75 | 54.05 | 58.84 | 59.96 | 58.98 | 65.35 | 100 | | |
| | *A. megaguti* | 29.69 | 37.30 | 31.15 | 57.26 | 60.00 | 62.15 | 61.84 | 67.12 | 67.88 | 100 | |
| | *A. dispar* | 28.40 | 36.03 | 29.01 | 54.85 | 58.89 | 60.41 | 59.58 | 65.38 | 83.65 | 69.16 | 100 |

**c** Pairwise nucleotide sequence similarity across *rfb* (sense vs. antisense) minioperons across ***P. distasonis* strains**

Doublet minioperons (column groups: - - BC: CBBP-1 to CL03T12C09; ++ CB: ATCC_8503 to FDAARGOS_615)

| | | CBBP-1 | CL03T12C09 | ATCC_8503 | 82G9 | NBRC_113806 | FDAARGOS_759 | CavFT-hAR46 | CavFT-hAR56 | FDAARGOS_615 |
|---|---|---|---|---|---|---|---|---|---|---|
| - - BC | CBBP-1 | 100 | | | | | | | | |
| | CL03T12C09 | 98.48 | 100 | | | | | | | |
| ++ CB | ATCC_8503 | 98.48 | 98.94 | 100 | | | | | | |
| | 82G9 | 98.53 | 99.00 | 99.82 | 100 | | | | | |
| | NBRC_113806 | 99.18 | 98.48 | 98.48 | 98.53 | 100 | | | | |
| | FDAARGOS_759 | 98.65 | 99.00 | 99.82 | 99.88 | 98.53 | 100 | | | |
| | CavFT-hAR46 | 98.65 | 99.00 | 99.82 | 99.88 | 98.53 | 100 | 100 | | |
| | CavFT-hAR56 | 98.65 | 99.00 | 99.82 | 99.88 | 98.53 | 100 | 100 | 100 | |
| | FDAARGOS_615 | 98.83 | 99.24 | 98.88 | 98.94 | 98.65 | 98.94 | 98.94 | 98.94 | 100 |

Triplet minioperons

| | | CavFT-hAR46 | CavFT-hAR56 | FDAARGOS_615 | FDAARGOS_759 | CL03T12C09 | CavFT-hAR46 | CavFT-hAR56 |
|---|---|---|---|---|---|---|---|---|
| - - - CGF | CavFT-hAR46 | 100 | | | | | | |
| | CavFT-hAR56 | 100 | 100 | | | | | |
| | FDAARGOS_615 | 98.52 | 98.52 | 100 | | | | |
| | FDAARGOS_759 | 99.26 | 99.26 | 97.90 | 100 | | | |
| +++ FGC | CL03T12C09 | 99.26 | 99.26 | 97.90 | 100 | 100 | | |
| | CavFT-hAR46 | 100 | 100 | 98.52 | 99.26 | 99.26 | 100 | |
| | CavFT-hAR56 | 100 | 100 | 98.52 | 99.26 | 99.26 | 100 | 100 |

Triplet minioperons

| | | NBRC_113806 | FDAARGOS_615 | 82G9 | FDAARGOS_615 | CBBP-1 |
|---|---|---|---|---|---|---|
| - - - DCA | NBRC_113806 | 100 | | | | |
| | FDAARGOS_615 | 99.95 | 100 | | | |
| +++ ACD | 82G9 | 87.82 | 87.86 | 100 | | |
| | FDAARGOS_615 | 99.95 | 100 | 87.86 | 100 | |
| | CBBP-1 | 99.83 | 99.79 | 87.74 | 99.79 | 100 |

**Supplementary Figure 3**. Sequence homology of *rfb* minioperons at the species and genus level, using **A)** *Parabacteroides* (12 genomes), **B)** *Alistipes* (11 genomes), and **C)** *Parabacteroides distasonis* (9 genomes) as models.

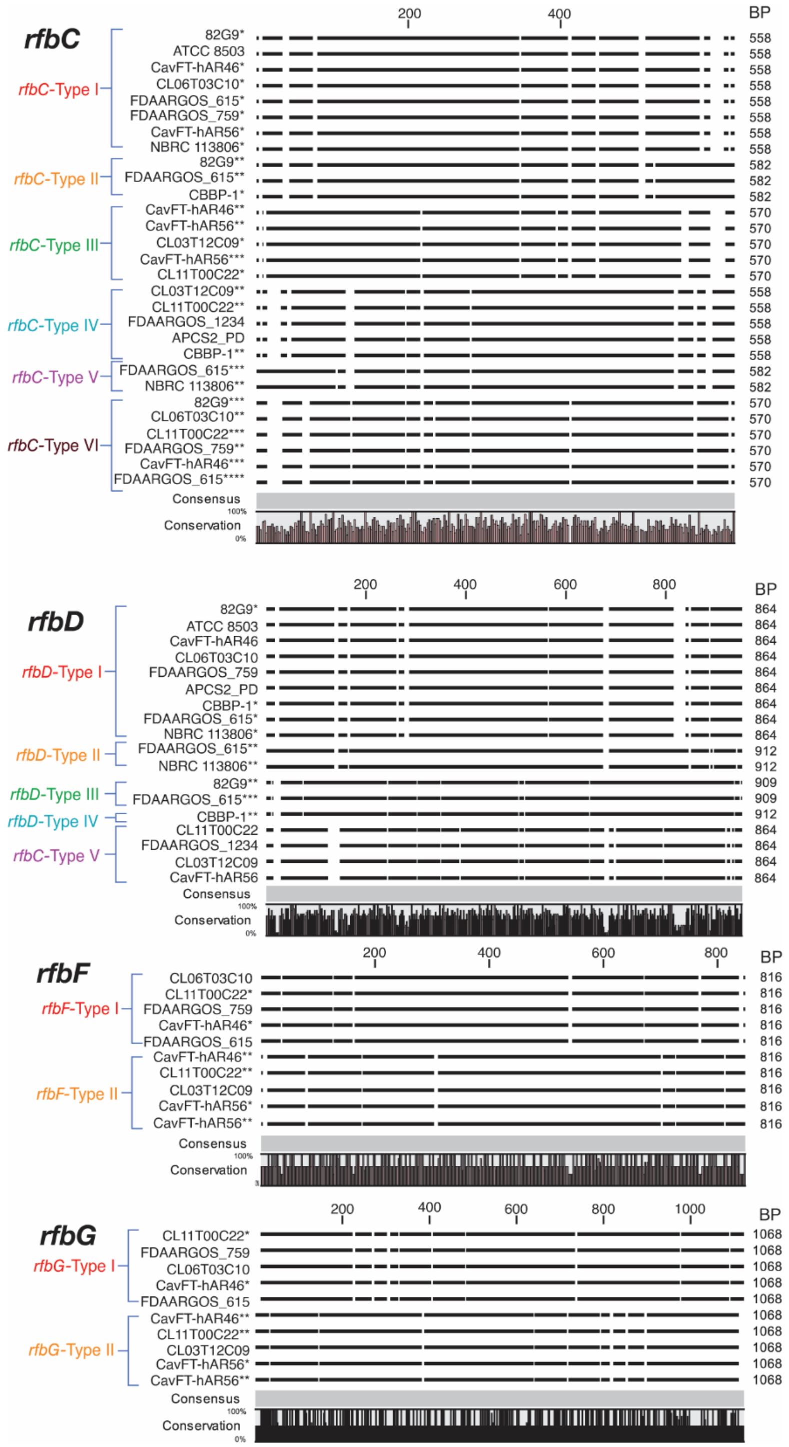

**Supplementary Figure 4.** Typing of the *rfbC*, *rfbD*, *rfbF*, and *rfbG* genes in 13 *Parabacteroides distasonis* strains utilizing the *rfbA*-Typing methodology (Bank et al., 2022).

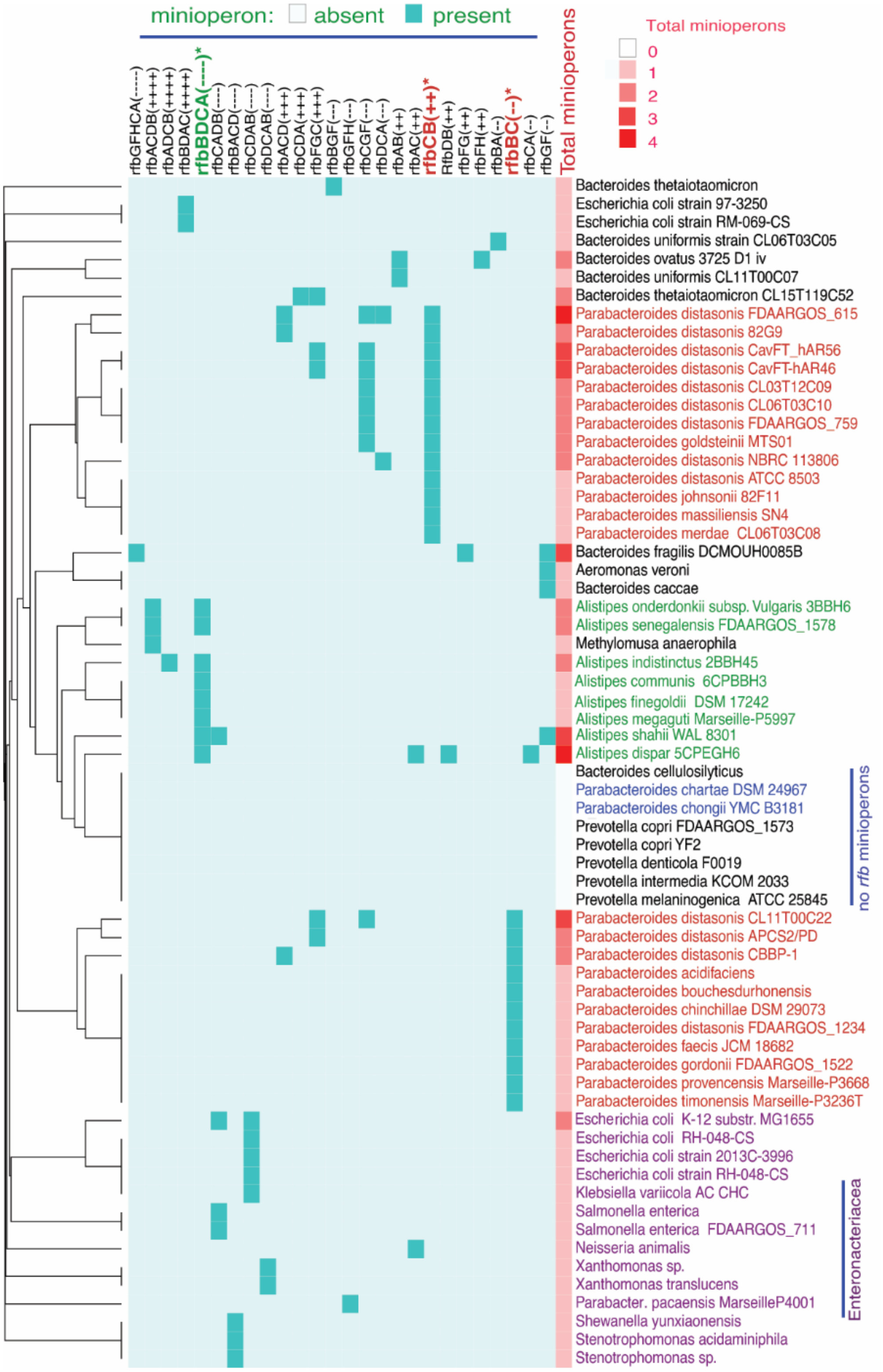

**Supplementary Figure 5**. Genera-wide phylogenetic analysis based on the *rfb* mini/operon content in species genomes.

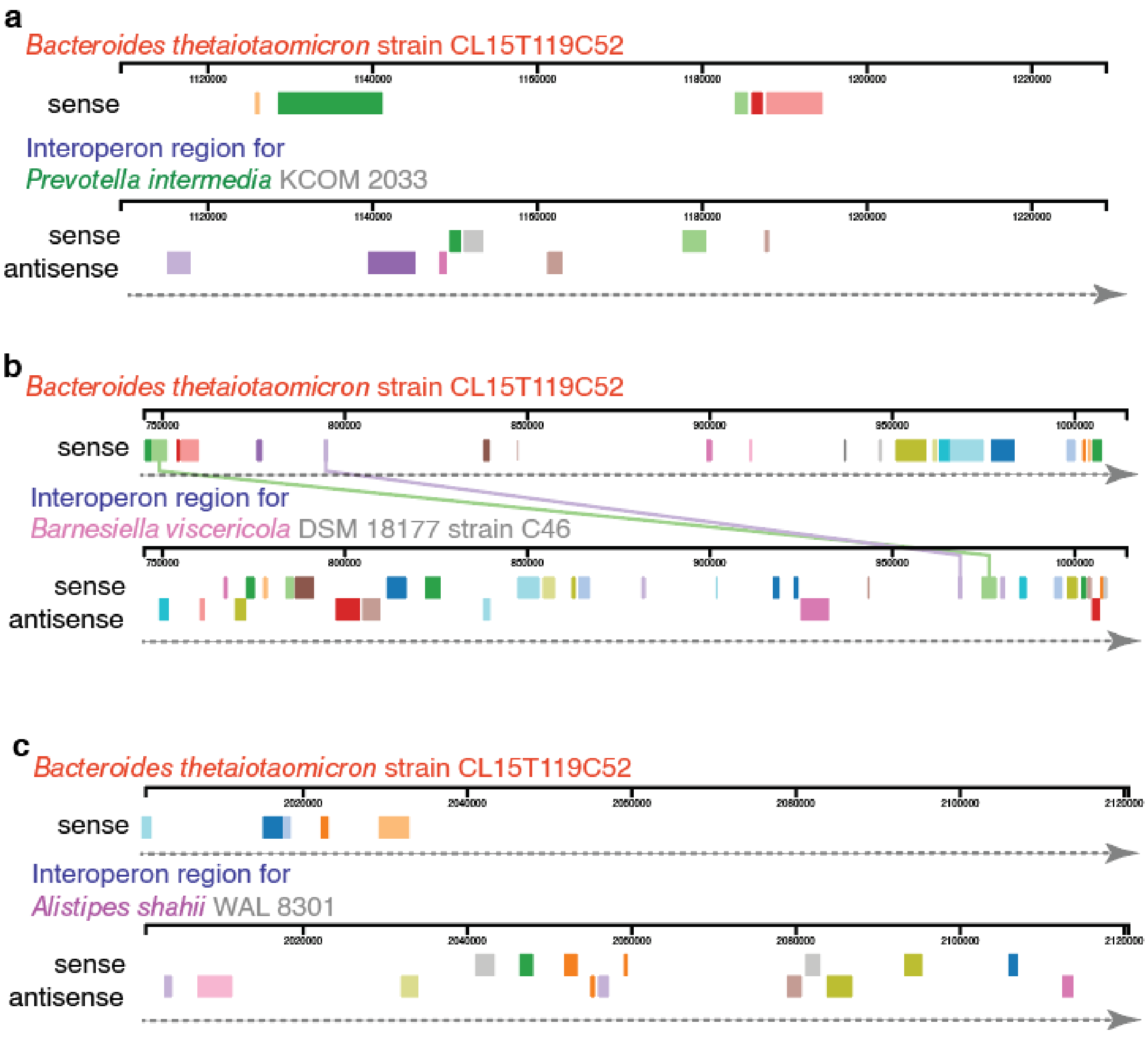

**Supplementary Figure 6**. Pairwise alignment of *B. thetaiotaomicron* genomes as source of inter-minioperon 'foreign DNA' within other *Bacteroidota*; **A)** *Prevotella intermedia*, 21% of inter-minioperon region matches *B. thetaiotaomicron* DNA, **B)** *Barnesiella viscericola*, 28%, and **C)** *Alistipes shahii*, 21%. Insertion details for these and other *Bacteroidota* are available in **Supplementary Table 5**.

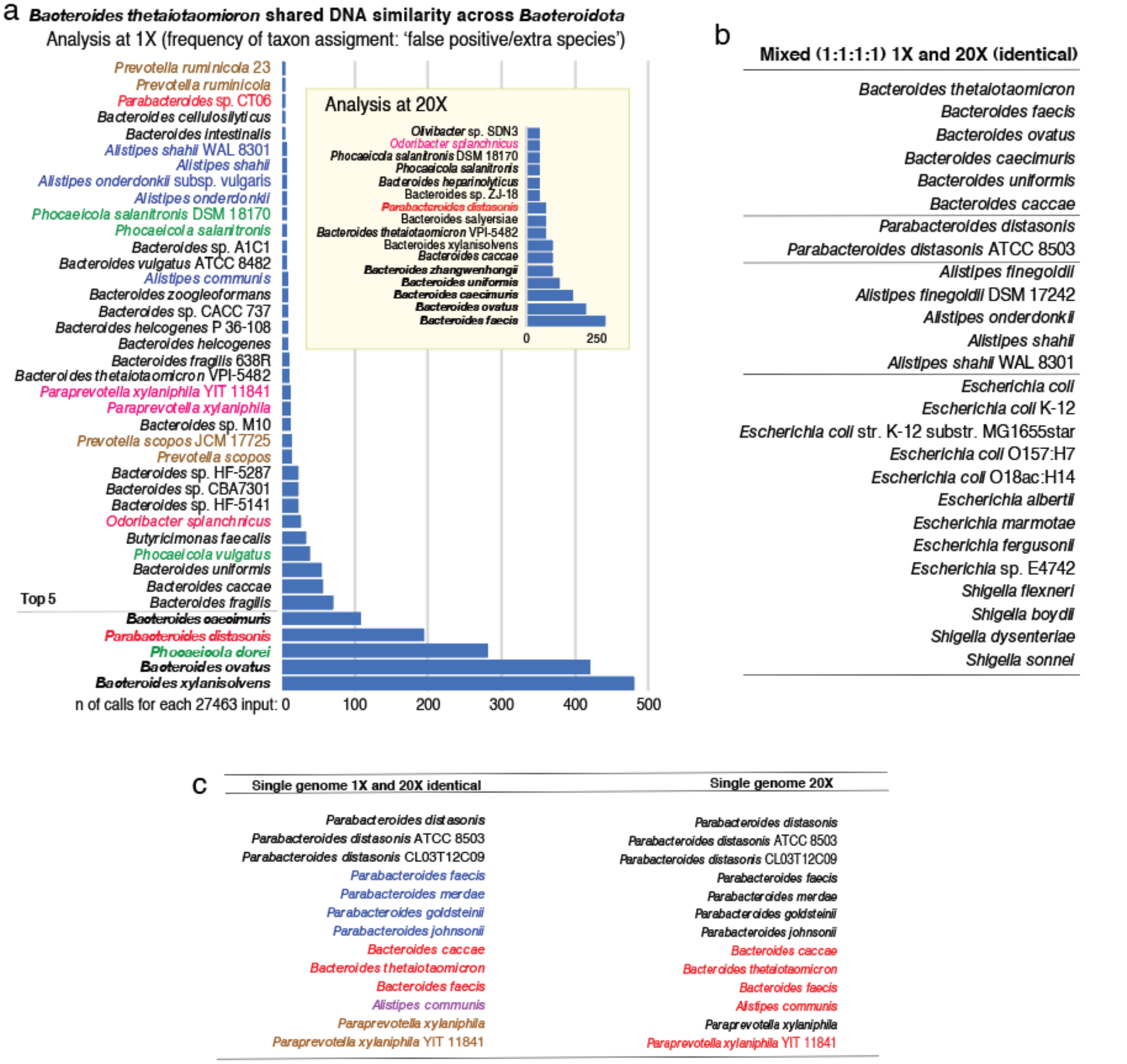

**Supplementary Figure 7**. Analysis of 'extra species' presence in a simulated metagenomics workflow for a community species identification (at 1X and 20X abundance). **A)** Frequency of 'extra species' called from *B. thetaiotaomicron* genome fragment. **B)** Top metagenomics results for simulated community analysis of mixed *B. thetaiotaomicron*, *P. distasonis*, *A. finegoldii*, and *E. coli* genomes. **C)** List of 'extra species' assigned by metagenomics of the *P. distasonis* genome fragmented and analyzed at 1X and 20X.

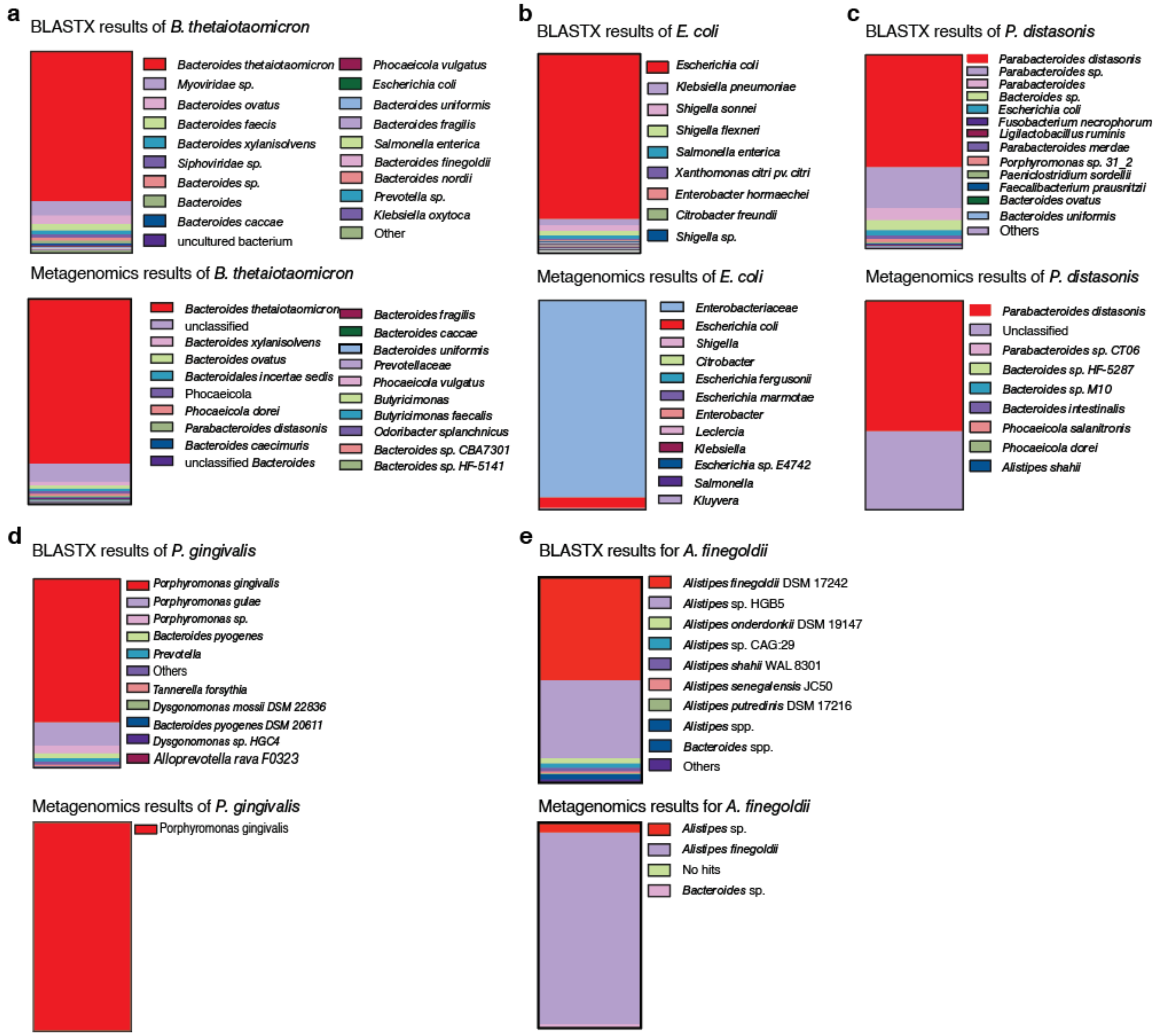

**Supplementary Figure 8**. BLAST and BLASTX performance for **A)** *B. thetaiotaomicron*, **B)** *E. coli*, **C)** *P. distasonis*, **D)** *P. gingivalis*, and **E)** *A. finegoldii*.

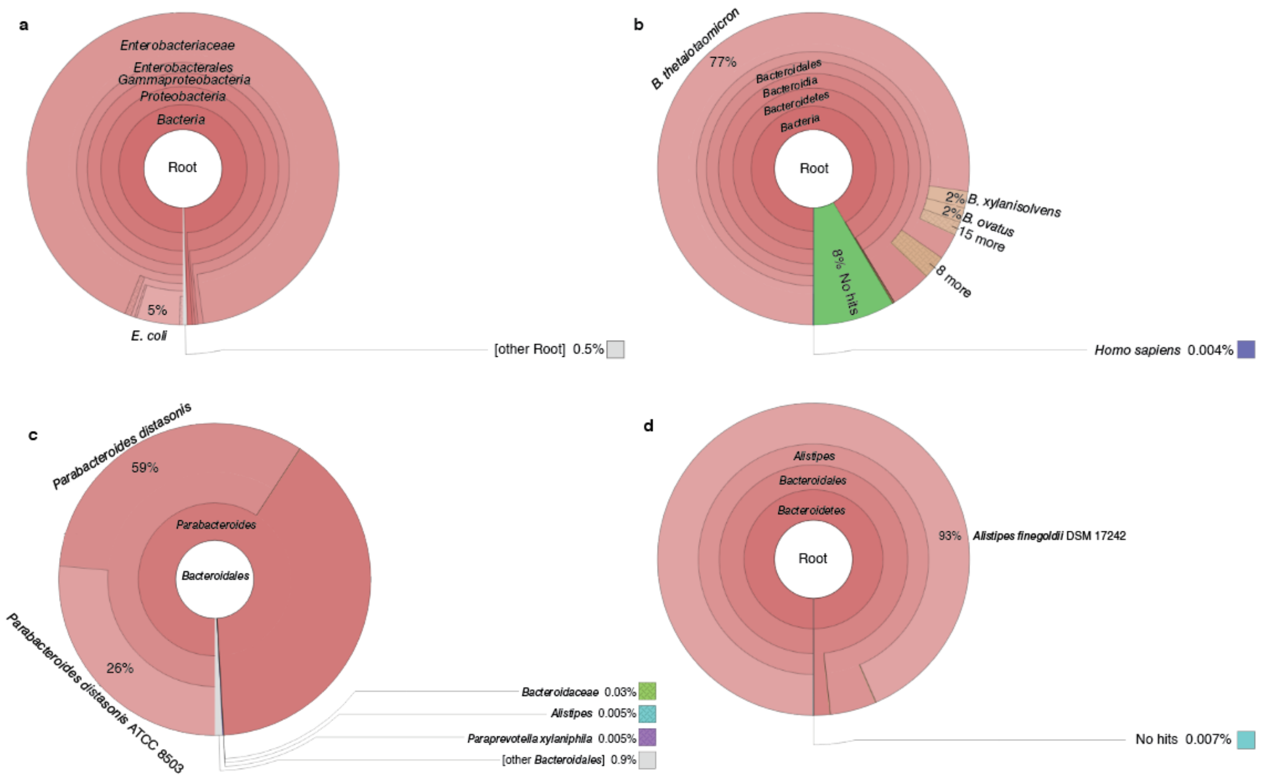

**Supplementary Figure 9**. BLAST and BLASTX results for induvial genome taxonomy of **A)** *Escherichia coli* K12, **B)** *Bacteroides thetaiotaomicron* CL15T119C52, **C)** *Parabacteroides distasonis* ATCC 8503, and **D)** *Alistipes finegoldii* DSM 17242. Whole genome sequences were split into 250bp inputs to simulate DNA-fragments reads. Analysis were conducted in various iterations, with 1x and 20x, and various fragment lengths (125bp to 16000 bp; not shown).

**Supplementary Table 1**. Top BLAST hits for *rfbACDB* and *rfbBDCA* operon primers developed for *Alistipes* spp.

| *Alistipes* operon | Primer sequence | BLAST Results | Max Score | Total Score | Query Cover | E value | Per. ident | Acc. Len | Accession |
|---|---|---|---|---|---|---|---|---|---|
| *rfbBDCA* | 5'-TCGATTCCCTCCTCGAACTGCAGC | Alistipes communis 6CPBBH3 DNA, complete genome | 40.1 | 40.1 | 100% | 2.4 | 95.83 | 3302129 | AP019739.1 |
| | | Alistipes dispar 5CPEGH6 DNA, complete genome | 40.1 | 80.3 | 100% | 2.4 | 95.83 | 2962376 | AP019736.1 |
| | | Alistipes communis 5CBH24 DNA, complete genome | 40.1 | 40.1 | 100% | 2.4 | 95.83 | 3301347 | AP019735.1 |
| | | Alistipes sp. Marseille-P5997 genome assembly, chromosome: contig00001 | 40.1 | 40.1 | 100% | 2.4 | 95.83 | 3270862 | LR027382.1 |
| | | Haemonchus contortus strain NZ_Hco_NP chromosome 2 | 38.2 | 74.3 | 79% | 9.6 | 100 | 77295023 | CP035801.1 |
| | | Alistipes onderdonkii subsp. vulgaris 5NYCFAH2 DNA, complete genome | 38.2 | 38.2 | 95% | 9.6 | 95.65 | 3312682 | AP019738.1 |
| | | Alistipes onderdonkii subsp. vulgaris 5CPYCFAH4 DNA, complete genome | 38.2 | 38.2 | 95% | 9.6 | 95.65 | 3312673 | AP019737.1 |
| | | Alistipes onderdonkii subsp. vulgaris 3BBH6 DNA, complete genome | 38.2 | 76.3 | 95% | 9.6 | 95.65 | 3507492 | AP019734.1 |
| | 3'-ACCTCNACGAAGATCGAGGCCTCGG | Alistipes onderdonkii subsp. vulgaris 5NYCFAH2 DNA, complete genome | 44.1 | 124 | 100% | 0.21 | 96 | 3312682 | AP019738.1 |
| | | Alistipes onderdonkii subsp. vulgaris 5CPYCFAH4 DNA, complete genome | 44.1 | 124 | 100% | 0.21 | 96 | 3312673 | AP019737.1 |
| | | Alistipes onderdonkii subsp. vulgaris 3BBH6 DNA, complete genome | 44.1 | 88.2 | 100% | 0.21 | 96 | 3507492 | AP019734.1 |
| | | Alistipes onderdonkii strain DSM 19147 chromosome, complete genome | 44.1 | 88.2 | 100% | 0.21 | 96 | 3869542 | CP102251.1 |
| | | uncultured Alistipes sp. isolate min17_bin03 genome assembly, chromosome: 1 | 44.1 | 44.1 | 100% | 0.21 | 96 | 2621094 | OV789733.1 |
| | | Alistipes finegoldii CE91-St15 DNA, complete genome | 44.1 | 88.2 | 100% | 0.21 | 96 | 4117255 | AP025581.1 |
| | | Alistipes onderdonkii isolate KR001_HAM_0061 chromosome, complete genome | 44.1 | 124 | 100% | 0.21 | 96 | 3395839 | CP107205.1 |
| | | Alistipes onderdonkii CE91-St18 DNA, complete genome | 44.1 | 124 | 100% | 0.21 | 96 | 3812179 | AP025562.1 |
| | | Alistipes finegoldii DSM 17242, complete genome | 44.1 | 80.3 | 100% | 0.21 | 96 | 3734239 | CP003274.1 |
| *rfbACDB* | 5'-AACTGCCTCTCGATCGAGGAGAA | Alistipes sp. Marseille-P5997 genome assembly, chromosome: contig00001 | 46.1 | 46.1 | 100% | 0.026 | 100 | 3270862 | LR027382.1 |
| | | Alistipes shahii WAL 8301 chromosome, complete genome | 46.1 | 78.3 | 100% | 0.026 | 100 | 3809618 | CP102253.1 |
| | | Alistipes senegalensis JC50 chromosome, complete genome | 46.1 | 78.3 | 100% | 0.026 | 100 | 4023197 | CP102252.1 |
| | | Alistipes shahii WAL 8301 draft genome | 46.1 | 78.3 | 100% | 0.026 | 100 | 3763317 | FP929032.1 |
| | | Bacteroidales bacterium isolate nC33_bin.4.fa chromosome | 46.1 | 46.1 | 100% | 0.026 | 100 | 1703659 | CP091721.1 |
| | | Alistipes senegalensis strain FDAARGOS_1578 chromosome, complete genome | 46.1 | 78.3 | 100% | 0.026 | 100 | 4023294 | CP085931.1 |
| | | uncultured Alistipes sp. isolate min17_bin03 genome assembly, chromosome: 1 | 42.1 | 74.3 | 91% | 0.41 | 100 | 2621094 | OV789733.1 |
| | | Anopheles stephensi strain Indian chromosome 2R | 40.1 | 74.3 | 86% | 1.6 | 100 | 58051481 | CP032299.1 |
| | | Anopheles stephensi strain SDA-500 chromosome 2R | 40.1 | 40.1 | 86% | 1.6 | 100 | 62568927 | CP032232.1 |
| | | Alistipes ihumii AP11 chromosome, complete genome | 40.1 | 120 | 86% | 1.6 | 100 | 2780015 | CP102294.1 |
| | | Anopheles maculipalpis genome assembly, chromosome: 2 | 40.1 | 40.1 | 86% | 1.6 | 100 | 98751411 | OX030895.1 |
| | | Mus musculus BAC clone RP24-194F23 from 5, complete sequence | 40.1 | 78.3 | 86% | 1.6 | 100 | 178113 | AC121917.2 |
| | | PREDICTED: Aquila chrysaetos chrysaetos FAT atypical cadherin 4 (FAT4), mRNA | 38.2 | 38.2 | 82% | 6.4 | 100 | 16978 | XM_030020759.1 |
| | | Alistipes onderdonkii subsp. vulgaris 5NYCFAH2 DNA, complete genome | 38.2 | 76.3 | 100% | 6.4 | 95.65 | 3312682 | AP019738.1 |
| | | Alistipes onderdonkii subsp. vulgaris 5CPYCFAH4 DNA, complete genome | 38.2 | 76.3 | 100% | 6.4 | 95.65 | 3312673 | AP019737.1 |
| | | Alistipes onderdonkii subsp. vulgaris 3BBH6 DNA, complete genome | 38.2 | 76.3 | 100% | 6.4 | 95.65 | 3507492 | AP019734.1 |
| | 3'-AACATCGACCTGATACGGGT | Alistipes onderdonkii subsp. vulgaris 5NYCFAH2 DNA, complete genome | 40.1 | 120 | 100% | 1.6 | 100 | 3312682 | AP019738.1 |
| | | Alistipes onderdonkii subsp. vulgaris 5CPYCFAH4 DNA, complete genome | 40.1 | 120 | 100% | 1.6 | 100 | 3312673 | AP019737.1 |
| | | Alistipes onderdonkii subsp. vulgaris 3BBH6 DNA, complete genome | 40.1 | 80.3 | 100% | 1.6 | 100 | 3507492 | AP019734.1 |
| | | Alistipes onderdonkii strain DSM 19147 chromosome, complete genome | 40.1 | 80.3 | 100% | 1.6 | 100 | 3869542 | CP102251.1 |
| | | Alistipes onderdonkii CE91-St18 DNA, complete genome | 40.1 | 120 | 100% | 1.6 | 100 | 3812179 | AP025562.1 |
| | | Alistipes senegalensis strain FDAARGOS_1578 chromosome, complete genome | 40.1 | 40.1 | 100% | 1.6 | 100 | 4023294 | CP085931.1 |
| | | Streptomyces nitrosporeus strain ATCC 12769 chromosome, complete genome | 40.1 | 40.1 | 100% | 1.6 | 100 | 7581562 | CP023702.1 |
| | | Alistipes onderdonkii isolate KR001_HAM_0061 chromosome, complete genome | 40.1 | 120 | 100% | 1.6 | 100 | 3395839 | CP107205.1 |
| | | Alistipes senegalensis JC50 chromosome, complete genome | 40.1 | 40.1 | 100% | 1.6 | 100 | 4023197 | CP102252.1 |
| | | Alistipes finegoldii CE91-St15 DNA, complete genome | 40.1 | 72.4 | 100% | 1.6 | 100 | 4117255 | AP025581.1 |
| | | Alistipes finegoldii DSM 17242, complete genome | 40.1 | 80.3 | 100% | 1.6 | 100 | 3734239 | CP003274.1 |

**Supplementary Table 2.** BLAST results indicate that *rfb* minioperon sequences in *P. distasonis* are only found in *P. distasonis*, with low coverage in some *Bacteroidota* and *Proteobacteria*.

| **Minioperon** | ***rfb* gene and sequence Type (I-VI)*** | | | | | | **Complete genome Blast hits bacterial kingdom lineage score for complete **** | | | | | |
|---|---|---|---|---|---|---|---|---|---|---|---|---|
| | **A** | **B** | **C** | **D** | **F** | **G** | ***Bacterial Species Strains*** | ***Pathogen Yes/No*** | ***Phylum*** | **Cover %** | **E value** | **Identity %** |
| *rfb* CB(++) | - | I | I | - | - | - | *P. distasonis* (15) | Human | Bacteroides | 100 | 0 | 100 - 98.8 |
| | | | | | | | *M. anaerophila** | No | Firmicutes | 3 | 5.00E-06 | 86.67 |
| *rfb* BC(--) | - | II | IV | - | - | - | *P. distasonis* (15) | Human | Bacteroides | 100 | 0 | 100 - 98.8 |
| | | | | | | | *M. anaerophila** | No | Firmicutes | 3 | 5.00E-06 | 86.67 |
| *rfb* ACD(+++) | III | - | II | III | - | - | *P. distasonis* (5) | Human | Bacteroides | **99** | 0 | 100-88 |
| | | | | & | | | *B. cellulosilyticus** | No | | 21 | 6.00E-56 | 75.1 |
| | | | | IV | | | *Stenotrophomonas** | Human | Proteobacteria | 2 | 9.00E-10 | 88.6 |
| | | | | | | | *Xanthomonas* (3) | Plant | Proteobacteria | 2 | 9.00E-10 | 91 - 89.6 |
| | | | | | | | *X. translucens* (18) | Plant | Proteobacteria | 2 | 4.00E-08 | 89.6 -85.7 |
| | | | | | | | *S. acidaminiphila* | Human | Proteobacteria | 2 | 9.00E-05 | 85.25 |
| | | | | | | | *S. yunxiaonensis** | Human | Proteobacteria | 1 | 1.00E-03 | 100 |
| | | | | | | | *N. animalis* (2) | Human | Proteobacteria | 2 | 1.50E-02 | 83.61 |
| *rfb* DCA(---) | IV | - | V | II | - | - | *P.distasonis* (5) | Human | Bacteroides * | 99 | 0 | 100 - 88 |
| | | | | | | | *B. cellulosilyticus** | No | | 21 | 6.00E-56 | 75.1 |
| | | | | | | | *Stenotrophomonas** | Human | Proteobacteria | 2 | 9.00E-10 | 88.6 |
| | | | | | | | *Xanthomonas* (3) | Plant | Proteobacteria | 2 | 9.00E-10 | 91.4 -89.7 |
| | | | | | | | *X. translucens* (18) | Plant | Proteobacteria | 2 | 4.00E-08 | 89.7 -85.7 |
| | | | | | | | *S. acidaminiphila** | Human | Proteobacteria | 2 | 9.00E-05 | 85.25 |
| | | | | | | | *S. yunxiaonensis** | Human | Proteobacteria | 1 | 1.00E-03 | 100 |
| | | | | | | | *N. animalis* (2) | Human | Proteobacteria | 2 | 1.50E-02 | 83.61 |
| *rfb* FGC(+++) | - | - | VI | - | II | II | *P. distasonis* (9) | Human | Bacteroides | 100 | 0 | 100 - 98.8 |
| | | | | | | | *B.thetaiotaomicron** | Human | | 24 | 1.00E-53 | 79.17 |
| | | | | | | | *B. caccae* (3) | Human | | 23 | 6.00E-47 | 79.2 -79.3 |
| | | | | | | | *A. veroni** | Human | Proteobacteria | 1 | 1.00E-05 | 87.72 |
| *rfb* CGF(---) | - | - | III | - | I | I | *P. distasonis* (9) | Human | Bacteroides | 100 | 0 | 100 - 98.8 |
| | | | | | | | *B. thetaiotaomicron** | Human | | 24 | 1.00E-53 | 79.2 |
| | | | | | | | *B. caccae* (3) | Human | | 23 | 6.00E-47 | 79.8 -79.3 |
| | | | | | | | A. veroni* | Human | Proteobacteria | 1 | 1.00E-05 | 87.72 |

**Rfb* -Type for each mini-operon *rfb* gene is also specified here. **Has only one strain in BLAST result

**Supplementary Table 3.** Summary inter-minioperon data for *rfbFGC -> rfbA, rfbCB -> rfbD,* and *fimCBA -> fimE* where present in three *P. distasonis* strains. * Denotes value could not be computed due to insufficient number of data points. See **Extended Data 1-3** for full inter-minioperon data.

| | **CavFT-hAR46 *rfbFGC* -> *rfbA* Interoperon Homology** | | | | **CavFT-hAR56 *rfbFGC* -> *rfbA* Interoperon Homology** | | | | **CL11T00C22 *rfbFGC* -> *rfbA* Interoperon Homology** | | | |
|---|---|---|---|---|---|---|---|---|---|---|---|---|
| | Total (bp) | Total (%) | Average size | ST.DEV size | Total (bp) | Total (%) | Average size | ST.DEV size | Total (bp) | Total (%) | Average size | ST.DEV size |
| *B. thetaiotamicron* | 37357 | 35.93021131 | 1009.648649 | 936.0901315 | 36862 | 35.45275307 | 1023.944444 | 945.0849378 | 73980 | 45.6308943 | 1155.9375 | 1342.003263 |
| *B. fragilis* | 35570 | 34.21146281 | 1111.5625 | 781.1221844 | 36288 | 34.90069728 | 1209.6 | 912.3378307 | 45843 | 28.27598118 | 955.0625 | 830.0099822 |
| *B. vulgatus* | 33725 | 32.43693 | 822.5609756 | 717.1880872 | 49216 | 47.3344554 | 1069.913043 | 1086.902261 | 57206 | 35.28468423 | 1100.115385 | 1838.520917 |
| *B. ovatus* | 25069 | 24.1115311 | 1044.541667 | 890.8435172 | 24619 | 23.67780717 | 1070.391304 | 880.9956526 | 69928 | 43.13161904 | 1043.701493 | 1185.202527 |
| *B. dorei* | 33487 | 32.20801954 | 1046.46875 | 832.4893096 | 33258 | 31.98653522 | 923.8333333 | 841.1978704 | 61709 | 38.06213647 | 1028.483333 | 1776.602752 |
| *A. communis* | 6197 | 5.960315857 | 885.2857143 | 700.6263184 | 6051 | 5.819668189 | 864.4285714 | 671.4873682 | 20335 | 12.54263633 | 884.1304348 | 573.7144764 |
| *A. finegoldii* | 9928 | 9.548816497 | 709.1428571 | 499.4188711 | 9830 | 9.45419572 | 756.1538462 | 489.5495968 | 17059 | 10.52199819 | 656.1153846 | 559.8838685 |
| *A. indistinctus* | 21323 | 20.50860336 | 852.92 | 837.4620847 | 20927 | 20.12695359 | 775.0740741 | 807.7140072 | 29227 | 18.02722557 | 789.9189189 | 664.5598534 |
| *A. onderdonkii* | 14332 | 13.78461302 | 895.75 | 569.6422854 | 13646 | 13.12430873 | 758.1111111 | 500.408761 | 25614 | 15.79872569 | 853.8 | 667.2685134 |
| *A. dispar* | 11237 | 10.80782141 | 1021.545455 | 650.6225271 | 10001 | 9.618658331 | 1000.1 | 743.3623985 | 15447 | 9.527715926 | 772.35 | 732.10089 |
| *E. coli* | 1815 | 1.745679084 | 453.75 | 258.1012398 | 1965 | 1.889877374 | 393 | 271.0498109 | 1803 | 1.112091138 | 450.75 | 261.0981616 |
| *E. albertii* | 1296 | 1.246501428 | 432 | 348.0387909 | 824 | 0.792498197 | 824 | * | 1791 | 1.104689534 | 255.8571429 | 263.5883334 |
| *E. fergusonii* | 1310 | 1.259966721 | 655 | 89.09545443 | 829 | 0.797307045 | 276.3333333 | 295.4950648 | 1480 | 0.912864606 | 493.3333333 | 195.3211031 |
| *E. hermannii* | 0 | 0 | 0 | 0 | 0 | 0 | * | * | 496 | 0.305933003 | 165.3333333 | 147.5545097 |
| *E. vulneris* | 753 | 0.724240413 | 376.5 | 55.86143571 | 753 | 0.724212551 | 376.5 | 55.86143571 | 722 | 0.445329896 | 361 | 33.9411255 |

| | **CavFT-hAR46 *rfbCB* -> *rfbD* Interoperon Homology** | | | | **CavFT-hAR56 *rfbCB* -> *rfbD* Interoperon Homology** | | | | **CL11T00C22 *fimCBA* -> *fimE* Interoperon Homology** | | | |
|---|---|---|---|---|---|---|---|---|---|---|---|---|
| | Total (bp) | Total (%) | Average size | ST.DEV size | Total (bp) | Total (%) | Average size | ST.DEV size | Total (bp) | Total (%) | Average size | ST.DEV size |
| *B. thetaiotamicron* | 266079 | 36.13804437 | 1051.695652 | 1033.362099 | 175383 | 23.81982643 | 958.3770492 | 1004.968446 | 254153 | 23.13162984 | 927.5656934 | 841.4554096 |
| *B. fragilis* | 271003 | 36.80680715 | 1066.940945 | 1024.1811 | 274405 | 37.2686034 | 980.0178571 | 971.4169138 | 287167 | 26.13638536 | 976.7585034 | 793.122743 |
| *B. vulgatus* | 266603 | 36.20921247 | 1079.364372 | 1096.898687 | 269060 | 36.54266661 | 1059.291339 | 1109.591511 | 288859 | 26.29038203 | 992.6426117 | 741.913883 |
| *B. ovatus* | 232757 | 31.6123512 | 977.9705882 | 1045.377296 | 225310 | 30.60071439 | 983.8864629 | 1000.978252 | 263475 | 23.98006781 | 893.1355932 | 785.1044659 |
| *B. dorei* | 254471 | 34.56148095 | 1078.266949 | 1031.194618 | 257516 | 34.97480612 | 1030.064 | 1031.768472 | 282589 | 25.71972059 | 1002.088652 | 745.5019394 |
| *A. communis* | 99977 | 13.57857351 | 854.5042735 | 747.0093203 | 46372 | 6.298061905 | 927.44 | 777.4270531 | 65154 | 5.929964277 | 868.72 | 740.0263794 |
| *A. finegoldii* | 104270 | 14.16163578 | 906.6956522 | 698.2327588 | 100931 | 13.70804982 | 877.6608696 | 701.2432679 | 90261 | 8.215067465 | 835.75 | 672.5103776 |
| *A. indistinctus* | 126524 | 17.1841067 | 832.3947368 | 653.1596882 | 128118 | 17.40048079 | 816.0382166 | 666.1495355 | 95721 | 8.712007099 | 811.1949153 | 754.7724541 |
| *A. onderdonkii* | 117651 | 15.9790027 | 840.3642857 | 728.9477704 | 120306 | 16.3394858 | 818.4081633 | 686.7970304 | 99179 | 9.026735534 | 944.5619048 | 783.26938 |
| *A. dispar* | 96652 | 13.12698208 | 840.4521739 | 731.5470419 | 90916 | 12.34785207 | 849.682243 | 749.2545931 | 88294 | 8.036041776 | 929.4105263 | 731.033755 |
| *E. coli* | 32103 | 4.360132286 | 594.5 | 431.2114899 | 14809 | 2.011299895 | 643.8695652 | 492.5015279 | 3825 | 0.348130788 | 546.4285714 | 575.6005146 |
| *E. albertii* | 29831 | 4.051556123 | 573.6730769 | 433.7715908 | 30612 | 4.157600945 | 518.8474576 | 407.3891562 | 3902 | 0.355138911 | 975.5 | 825.642578 |
| *E. fergusonii* | 33789 | 4.589119702 | 553.9180328 | 436.4353826 | 33430 | 4.540330576 | 576.3793103 | 443.611883 | 6668 | 0.606885253 | 833.5 | 720.9507611 |
| *E. hermannii* | 24430 | 3.318008652 | 508.9583333 | 444.5696919 | 23344 | 3.170489888 | 542.8837209 | 418.1507594 | 7687 | 0.699629116 | 512.4666667 | 439.3563257 |
| *E. vulneris* | 30100 | 4.088090889 | 578.8461538 | 509.903735 | 30030 | 4.078556004 | 536.25 | 457.3245217 | 6380 | 0.580673053 | 490.7692308 | 683.7359571 |

**Supplementary Table 4**. Ribosomal RNA (*rrn*) loci in other *Bacteroidota*. Note that they are similarly contiguous as observed *in P. distasonis*, likely indicating that *B. thetaiotaomicron* DNA insertions which disrupt these essential operons compromise bacterial survival and are thereby selected against.

| Genome ID | Genome Name | Accession | Annotation | Feature Ty | Start | End | Strand | Length (NA) | Product |
|---|---|---|---|---|---|---|---|---|---|
| 28131.18 | Prevotella intermedia strain KCOM 2033 | CP024696 | RefSeq | rRNA | 879563 | 879675 | - | 113 | 5S ribosomal RNA |
| 28131.18 | Prevotella intermedia strain KCOM 2033 | CP024696 | RefSeq | rRNA | 879786 | 882693 | - | 2908 | 23S ribosomal RNA |
| 28131.18 | Prevotella intermedia strain KCOM 2033 | CP024696 | RefSeq | rRNA | 883203 | 884742 | - | 1540 | 16S ribosomal RNA |
| 28131.18 | Prevotella intermedia strain KCOM 2033 | CP024696 | RefSeq | rRNA | 1491929 | 1492041 | - | 113 | 5S ribosomal RNA |
| 28131.18 | Prevotella intermedia strain KCOM 2033 | CP024696 | RefSeq | rRNA | 1492152 | 1495059 | - | 2908 | 23S ribosomal RNA |
| 28131.18 | Prevotella intermedia strain KCOM 2033 | CP024696 | RefSeq | rRNA | 1495569 | 1497108 | - | 1540 | 16S ribosomal RNA |
| 28131.18 | Prevotella intermedia strain KCOM 2033 | CP024696 | RefSeq | rRNA | 1757168 | 1757280 | - | 113 | 5S ribosomal RNA |
| 28131.18 | Prevotella intermedia strain KCOM 2033 | CP024696 | RefSeq | rRNA | 1757391 | 1760299 | - | 2909 | 23S ribosomal RNA |
| 28131.18 | Prevotella intermedia strain KCOM 2033 | CP024696 | RefSeq | rRNA | 1760648 | 1762187 | - | 1540 | 16S ribosomal RNA |
| 28131.18 | Prevotella intermedia strain KCOM 2033 | CP024696 | RefSeq | rRNA | 2105961 | 2106073 | + | 113 | 5S ribosomal RNA |
| 28131.18 | Prevotella intermedia strain KCOM 2033 | CP024696 | RefSeq | rRNA | 2220859 | 2222398 | + | 1540 | 16S ribosomal RNA |
| 28131.18 | Prevotella intermedia strain KCOM 2033 | CP024696 | RefSeq | rRNA | 2222747 | 2225655 | + | 2909 | 23S ribosomal RNA |
| 28131.18 | Prevotella intermedia strain KCOM 2033 | CP024696 | RefSeq | rRNA | 2225766 | 2225878 | + | 113 | 5S ribosomal RNA |

| Genome ID | Genome Name | Accession | Annotation | Feature Ty | Start | End | Strand | Length (NA) | Product |
|---|---|---|---|---|---|---|---|---|---|
| 818.1896 | Bacteroides thetaiotaomicron CL15T119C52 | CP075195 | RefSeq | rRNA | 1861265 | 1861371 | - | 107 | 5S ribosomal RNA |
| 818.1896 | Bacteroides thetaiotaomicron CL15T119C52 | CP075195 | RefSeq | rRNA | 1861550 | 1864420 | - | 2871 | 23S ribosomal RNA |
| 818.1896 | Bacteroides thetaiotaomicron CL15T119C52 | CP075195 | RefSeq | rRNA | 1864951 | 1866476 | - | 1526 | 16S ribosomal RNA |
| 818.1896 | Bacteroides thetaiotaomicron CL15T119C52 | CP075195 | RefSeq | rRNA | 3654432 | 3655957 | + | 1526 | 16S ribosomal RNA |
| 818.1896 | Bacteroides thetaiotaomicron CL15T119C52 | CP075195 | RefSeq | rRNA | 3656488 | 3659358 | + | 2871 | 23S ribosomal RNA |
| 818.1896 | Bacteroides thetaiotaomicron CL15T119C52 | CP075195 | RefSeq | rRNA | 3659537 | 3659643 | + | 107 | 5S ribosomal RNA |
| 818.1896 | Bacteroides thetaiotaomicron CL15T119C52 | CP075195 | RefSeq | rRNA | 3818471 | 3819990 | + | 1520 | 16S ribosomal RNA |
| 818.1896 | Bacteroides thetaiotaomicron CL15T119C52 | CP075195 | RefSeq | rRNA | 3820521 | 3823391 | + | 2871 | 23S ribosomal RNA |
| 818.1896 | Bacteroides thetaiotaomicron CL15T119C52 | CP075195 | RefSeq | rRNA | 3823570 | 3823676 | + | 107 | 5S ribosomal RNA |
| 818.1896 | Bacteroides thetaiotaomicron CL15T119C52 | CP075195 | RefSeq | rRNA | 4017998 | 4018087 | - | 90 | 5S ribosomal RNA |
| 818.1896 | Bacteroides thetaiotaomicron CL15T119C52 | CP075195 | RefSeq | rRNA | 4475271 | 4476796 | + | 1526 | 16S ribosomal RNA |
| 818.1896 | Bacteroides thetaiotaomicron CL15T119C52 | CP075195 | RefSeq | rRNA | 4477327 | 4480197 | + | 2871 | 23S ribosomal RNA |
| 818.1896 | Bacteroides thetaiotaomicron CL15T119C52 | CP075195 | RefSeq | rRNA | 4480376 | 4480482 | + | 107 | 5S ribosomal RNA |
| 818.1896 | Bacteroides thetaiotaomicron CL15T119C52 | CP075195 | RefSeq | rRNA | 5328411 | 5329936 | + | 1526 | 16S ribosomal RNA |
| 818.1896 | Bacteroides thetaiotaomicron CL15T119C52 | CP075195 | RefSeq | rRNA | 5330467 | 5333337 | + | 2871 | 23S ribosomal RNA |
| 818.1896 | Bacteroides thetaiotaomicron CL15T119C52 | CP075195 | RefSeq | rRNA | 5333516 | 5333622 | + | 107 | 5S ribosomal RNA |

| Genome ID | Genome Name | Accession | Annotation | Feature Ty | Start | End | Strand | Length (NA) | Product |
|---|---|---|---|---|---|---|---|---|---|
| 817.56 | Bacteroides fragilis strain DCMOUH0085B | JPHP01000002 | RefSeq | rRNA | 1 | 53 | - | 53 | 16S ribosomal RNA |
| 817.56 | Bacteroides fragilis strain DCMOUH0085B | JPHP01000031 | RefSeq | rRNA | 1 | 88 | + | 88 | 23S ribosomal RNA |
| 817.56 | Bacteroides fragilis strain DCMOUH0085B | JPHP01000053 | RefSeq | rRNA | 1 | 110 | - | 110 | 16S ribosomal RNA |
| 817.56 | Bacteroides fragilis strain DCMOUH0085B | JPHP01000143 | RefSeq | rRNA | 1 | 88 | + | 88 | 23S ribosomal RNA |
| 817.56 | Bacteroides fragilis strain DCMOUH0085B | JPHP01000100 | RefSeq | rRNA | 18 | 1557 | + | 1540 | 16S ribosomal RNA |
| 817.56 | Bacteroides fragilis strain DCMOUH0085B | JPHP01000076 | RefSeq | rRNA | 155 | 206 | + | 52 | 16S ribosomal RNA |
| 817.56 | Bacteroides fragilis strain DCMOUH0085B | JPHP01000031 | RefSeq | rRNA | 177 | 287 | + | 111 | 5S ribosomal RNA |
| 817.56 | Bacteroides fragilis strain DCMOUH0085B | JPHP01000143 | RefSeq | rRNA | 177 | 287 | + | 111 | 5S ribosomal RNA |
| 817.56 | Bacteroides fragilis strain DCMOUH0085B | JPHP01000100 | RefSeq | rRNA | 2032 | 4925 | + | 2894 | 23S ribosomal RNA |
| 817.56 | Bacteroides fragilis strain DCMOUH0085B | JPHP01000036 | RefSeq | rRNA | 6500 | 6608 | - | 109 | 5S ribosomal RNA |
| 817.56 | Bacteroides fragilis strain DCMOUH0085B | JPHP01000138 | RefSeq | rRNA | 43260 | 43370 | - | 111 | 5S ribosomal RNA |
| 817.56 | Bacteroides fragilis strain DCMOUH0085B | JPHP01000138 | RefSeq | rRNA | 43459 | 43546 | - | 88 | 23S ribosomal RNA |
| 817.56 | Bacteroides fragilis strain DCMOUH0085B | JPHP01000132 | RefSeq | rRNA | 51118 | 51215 | - | 98 | 5S ribosomal RNA |
| 817.56 | Bacteroides fragilis strain DCMOUH0085B | JPHP01000110 | RefSeq | rRNA | 366259 | 366369 | - | 111 | 5S ribosomal RNA |
| 817.56 | Bacteroides fragilis strain DCMOUH0085B | JPHP01000110 | RefSeq | rRNA | 366458 | 366545 | - | 88 | 23S ribosomal RNA |
| 817.56 | Bacteroides fragilis strain DCMOUH0085B | JPHP01000031 | RefSeq | rRNA | 456863 | 456972 | + | 110 | 16S ribosomal RNA |

| Genome ID | Genome Name | Accession | Annotation | Feature Ty | Start | End | Strand | Length (NA) | Product |
|---|---|---|---|---|---|---|---|---|---|
| 679935.3 | Alistipes finegoldii DSM 17242 | CP003274 | RefSeq | rRNA | 115704 | 115817 | - | 114 | 5S ribosomal RNA |
| 679935.3 | Alistipes finegoldii DSM 17242 | CP003274 | RefSeq | rRNA | 116001 | 118868 | - | 2868 | 23S ribosomal RNA |
| 679935.3 | Alistipes finegoldii DSM 17242 | CP003274 | RefSeq | rRNA | 119147 | 120561 | - | 1415 | 16S ribosomal RNA |
| 679935.3 | Alistipes finegoldii DSM 17242 | CP003274 | RefSeq | rRNA | 2181216 | 2182629 | + | 1414 | 16S ribosomal RNA |
| 679935.3 | Alistipes finegoldii DSM 17242 | CP003274 | RefSeq | rRNA | 2183071 | 2185938 | + | 2868 | 23S ribosomal RNA |
| 679935.3 | Alistipes finegoldii DSM 17242 | CP003274 | RefSeq | rRNA | 2186122 | 2186235 | + | 114 | 5S ribosomal RNA |

| Genome ID | Genome Name | Accession | Annotation | Feature Ty | Start | End | Strand | Length (NA) | Product |
|---|---|---|---|---|---|---|---|---|---|
| 837.126 | Porphyromonas gingivalis strain KCOM 2805 | CP024594 | RefSeq | rRNA | 285996 | 287537 | + | 1542 | 16S ribosomal RNA |
| 837.126 | Porphyromonas gingivalis strain KCOM 2805 | CP024594 | RefSeq | rRNA | 288288 | 291184 | + | 2897 | 23S ribosomal RNA |
| 837.126 | Porphyromonas gingivalis strain KCOM 2805 | CP024594 | RefSeq | rRNA | 291275 | 291385 | + | 111 | 5S ribosomal RNA |
| 837.126 | Porphyromonas gingivalis strain KCOM 2805 | CP024594 | RefSeq | rRNA | 364756 | 366297 | + | 1542 | 16S ribosomal RNA |
| 837.126 | Porphyromonas gingivalis strain KCOM 2805 | CP024594 | RefSeq | rRNA | 367047 | 369943 | + | 2897 | 23S ribosomal RNA |
| 837.126 | Porphyromonas gingivalis strain KCOM 2805 | CP024594 | RefSeq | rRNA | 370034 | 370144 | + | 111 | 5S ribosomal RNA |
| 837.126 | Porphyromonas gingivalis strain KCOM 2805 | CP024594 | RefSeq | rRNA | 633568 | 635109 | + | 1542 | 16S ribosomal RNA |
| 837.126 | Porphyromonas gingivalis strain KCOM 2805 | CP024594 | RefSeq | rRNA | 635860 | 638756 | + | 2897 | 23S ribosomal RNA |
| 837.126 | Porphyromonas gingivalis strain KCOM 2805 | CP024594 | RefSeq | rRNA | 638847 | 638957 | + | 111 | 5S ribosomal RNA |
| 837.126 | Porphyromonas gingivalis strain KCOM 2805 | CP024594 | RefSeq | rRNA | 1133092 | 1134633 | + | 1542 | 16S ribosomal RNA |
| 837.126 | Porphyromonas gingivalis strain KCOM 2805 | CP024594 | RefSeq | rRNA | 1135384 | 1138280 | + | 2897 | 23S ribosomal RNA |
| 837.126 | Porphyromonas gingivalis strain KCOM 2805 | CP024594 | RefSeq | rRNA | 1138371 | 1138481 | + | 111 | 5S ribosomal RNA |

**Supplementary Table 5**. Total and % DNA from select inter-minioperon regions of various *Bacteroidota* matching *B. thetaiotaomicron* DNA.

| | *B. fragilis* | *P. intermedia* | *B. viscericola* | *A. shahii* |
|---|---|---|---|---|
| Inter-minioperon distance (bp) | 236301 | 76877 | 310556 | 33052 |
| Total inter-minioperon space matching *B. thetaiotaomicron* DNA (bp) | 112035 | 16163 | 86235 | 6997 |
| % Inter-minioperon space matching *B. thetaiotaomicron* DNA | **47.4** | **21.0** | **27.8** | **21.2** |

**Supplementary Table 6**. Top 20 BLAST results of inter-minioperon DNA sequences from *B. thetaiotaomicron* and *B. fragilis* demonstrating the high-similarity matches which could lead to distorted metagenomic interpretations.

| Organism | Accession | *rfbFGC* end | *rfbA* start | Description | Max Score | Total Score | Query Cover | E value | Per. ident | Acc. Len | Accession |
|---|---|---|---|---|---|---|---|---|---|---|---|
| *B. thetaiotaomicron* CL15T119C52 | CP075195.1 | 1657979 | 1661329 | Bacteroides thetaiotaomicron strain CL15T119C52 chromosome, complete genome | 6189 | 6189 | 1 | 0 | 100 | 6865713 | CP075195.1 |
| | | | | Bacteroides thetaiotaomicron strain CL15T12C11 chromosome, complete genome | 6189 | 6189 | 1 | 0 | 100 | 6844923 | CP075193.1 |
| | | | | Bacteroides thetaiotaomicron strain BFG-510 chromosome, complete genome | 6139 | 6139 | 1 | 0 | 99.73 | 6573230 | CP103118.1 |
| | | | | Bacteroides thetaiotaomicron strain BFG-109 chromosome, complete genome | 6128 | 6128 | 1 | 0 | 99.67 | 5957022 | CP103077.1 |
| | | | | Bacteroides thetaiotaomicron strain VPI-BTDOT2 chromosome | 6106 | 6106 | 1 | 0 | 99.55 | 6467639 | CP083681.1 |
| | | | | Bacteroides thetaiotaomicron strain BFG-169 chromosome, complete genome | 6106 | 6106 | 1 | 0 | 99.55 | 6057350 | CP103072.1 |
| | | | | Bacteroides thetaiotaomicron strain BFG-129 chromosome, complete genome | 6106 | 6106 | 1 | 0 | 99.55 | 6282330 | CP103075.1 |
| | | | | Bacteroides thetaiotaomicron strain CL06T03C18 chromosome, complete genome | 6106 | 6106 | 1 | 0 | 99.55 | 5830764 | CP072242.1 |
| | | | | Bacteroides thetaiotaomicron strain BFG-498 chromosome, complete genome | 6100 | 6100 | 1 | 0 | 99.52 | 6103379 | CP103214.1 |
| | | | | Bacteroides thetaiotaomicron strain CL11T00C24 chromosome, complete genome | 6089 | 6089 | 1 | 0 | 99.46 | 6712089 | CP072224.1 |
| | | | | Bacteroides thetaiotaomicron F9-2 DNA, nearly complete genome | 6061 | 6061 | 1 | 0 | 99.31 | 6224560 | AP022660.1 |
| | | | | Bacteroides thetaiotaomicron strain BFG-119 chromosome, complete genome | 6061 | 6061 | 1 | 0 | 99.31 | 6246733 | CP103283.1 |
| | | | | Bacteroides thetaiotaomicron strain BFG-148 chromosome, complete genome | 6050 | 6050 | 1 | 0 | 99.25 | 6296712 | CP103095.1 |
| | | | | Bacteroides thetaiotaomicron strain VPI-2808B chromosome | 6039 | 6039 | 1 | 0 | 99.19 | 6225667 | CP083687.1 |
| | | | | Bacteroides thetaiotaomicron strain VPI-3443 chromosome | 6039 | 6039 | 1 | 0 | 99.19 | 6301124 | CP083685.1 |
| | | | | Bacteroides thetaiotaomicron strain VPI-5482A chromosome | 6039 | 6039 | 1 | 0 | 99.19 | 6254757 | CP083684.1 |
| | | | | Bacteroides thetaiotaomicron strain VPI-BT7853 chromosome | 6039 | 6039 | 1 | 0 | 99.19 | 6319080 | CP083683.1 |
| | | | | Bacteroides thetaiotaomicron strain BFG-576 chromosome, complete genome | 6039 | 6039 | 1 | 0 | 99.19 | 6558000 | CP103155.1 |
| | | | | Bacteroides thetaiotaomicron strain BFG-484 chromosome, complete genome | 6039 | 6039 | 1 | 0 | 99.19 | 6071302 | CP103153.1 |
| | | | | Bacteroides thetaiotaomicron strain BFG-446 chromosome, complete genome | 6039 | 6039 | 1 | 0 | 99.19 | 6465642 | CP103104.1 |
| Organism | Accession | *rfbD* end | *rfbAFG* start | Bacteroides ovatus strain 3725 D1 iv chromosome, complete genome | 6307 | 6307 | 1 | 0 | 100 | 7079631 | CP041395.1 |
| *B. fragilis* DCMOUH0085B | CP037440.1 | 1658013 | 1661427 | Bacteroides fragilis strain DCMOUH0085B chromosome, complete genome | 6307 | 6307 | 1 | 0 | 100 | 5504076 | CP037440.1 |
| | | | | Bacteroides fragilis strain IHMA_4 chromosome, complete genome | 6307 | 11156 | 1 | 0 | 100 | 5540606 | CP080295.1 |
| | | | | Bacteroides xylanisolvens strain APCS1/XY chromosome, complete genome | 5380 | 5380 | 0.85 | 0 | 100 | 6461058 | CP042282.1 |
| | | | | Bacteroides ovatus isolate KR001_HAM_0001 chromosome, complete genome | 4854 | 4854 | 1 | 0 | 92.33 | 6770402 | CP107192.1 |
| | | | | Alistipes finegoldii CE91-St15 DNA, complete genome | 4854 | 4854 | 1 | 0 | 92.33 | 4117255 | AP025581.1 |
| | | | | Odoribacteraceae bacterium CE91-St21 DNA, complete genome | 4854 | 4854 | 1 | 0 | 92.33 | 5743229 | AP025578.1 |
| | | | | Phocaeicola vulgatus MG01-10 DNA, complete genome | 4854 | 4854 | 1 | 0 | 92.33 | 4957189 | AP025240.1 |
| | | | | Phocaeicola vulgatus MG01-07 DNA, complete genome | 4854 | 4854 | 1 | 0 | 92.33 | 4985319 | AP025235.1 |
| | | | | Phocaeicola vulgatus MG01-03 DNA, complete genome | 4854 | 4854 | 1 | 0 | 92.33 | 5073285 | AP025232.1 |
| | | | | Bacteroides uniformis strain JCM13288 chromosome, complete genome | 4854 | 6707 | 1 | 0 | 92.33 | 5068087 | CP054204.1 |
| | | | | Bacteroides uniformis NBRC 113350 DNA, complete genome | 4848 | 4848 | 1 | 0 | 92.3 | 4734883 | AP019724.1 |
| | | | | Bacteroides fragilis strain DCMOUH0018B chromosome, complete genome | 4848 | 4848 | 1 | 0 | 92.31 | 5302644 | CP036542.1 |
| | | | | Bacteroides sp. PHL 2737 chromosome, complete genome | 4848 | 4848 | 1 | 0 | 92.31 | 5522430 | CP040630.1 |
| | | | | Bacteroides dorei isolate MGYG-HGUT-02478 genome assembly, chromosome: 1 | 4691 | 4691 | 0.99 | 0 | 91.47 | 5444912 | LR699004.1 |
| | | | | Bacteroides faecis strain BFG-493 chromosome, complete genome | 4691 | 4691 | 0.99 | 0 | 91.47 | 6287716 | CP103125.1 |
| | | | | Bacteroides thetaiotaomicron strain BFG-129 chromosome, complete genome | 2159 | 2159 | 0.89 | 0 | 79.6 | 6282330 | CP103075.1 |
| | | | | Parabacteroides johnsonii DSM 18315 chromosome, complete genome | 2121 | 2121 | 0.91 | 0 | 79.11 | 4749455 | CP102285.1 |
| | | | | Parabacteroides johnsonii strain FDAARGOS_1580 chromosome, complete genome | 2121 | 2121 | 0.91 | 0 | 79.11 | 4766157 | CP085975.1 |
| | | | | Bacteroides thetaiotaomicron strain BFG-576 chromosome, complete genome | 2002 | 3582 | 0.99 | 0 | 77.43 | 6558000 | CP103155.1 |

**Supplementary Table 7**. Gene counts of KEGG Pathways for select *Enterobacteriaceae* and *Bacteroidota*.

| | *K. variicola* | *E. coli* | *S. enterica* | *P. distasonis* | *B. thetaiotaomicron* | *B. fragilis* | *O. splanchnicus* | *A. finegoldii* | *P. melaninogenica* |
|---|---|---|---|---|---|---|---|---|---|
| **Metabolism** | **2143** | **1727** | **1714** | **1240** | **1303** | **1231** | **1045** | **913** | **787** |
| **Genetic Information Processing** | **369** | **358** | **364** | **314** | **297** | **302** | **288** | **260** | **263** |
| **Environmental Information Processing** | **559** | **461** | **495** | **167** | **126** | **127** | **176** | **125** | **80** |
| **Cellular Processes** | **140** | **221** | **157** | **62** | **52** | **54** | **61** | **54** | **56** |
| **Organismal Systems** | **91** | **91** | **91** | **91** | **91** | **91** | **91** | **91** | **91** |
| **Human Diseases** | **166** | **157** | **200** | **145** | **132** | **134** | **135** | **137** | **120** |
| **Brite Hierarchies** | **2471** | **2301** | **2252** | **1349** | **1180** | **1206** | **1190** | **1001** | **909** |
| **Not Included in Pathway or Brite** | **569** | **626** | **574** | **261** | **266** | **210** | **217** | **217** | **148** |
| **Grand Total** | **6508** | **5942** | **5847** | **3629** | **3447** | **3355** | **3203** | **2798** | **2454** |
